# From phenotype to receptor: validating physiological clustering of *Escherichia coli* phages through comprehensive receptor analysis

**DOI:** 10.1101/2024.11.29.626071

**Authors:** Tomoyoshi Kaneko, Toshifumi Osaka, Minoru Inagaki, Kento Habe, Takuma Okabe, Satoshi Tsuneda

**Author notes:** Address correspondence corresponding author to Satoshi Tsuneda, TWIns, 2-2 Wakamatsu-cho, Shinjuku-ku, Tokyo, Japan, 162-8480. **Authorship Attribution:** Tomoyoshi Kaneko: Conceptualization, Data curation, Formal analysis, Writing - original draft, Investigation Osaka Toshifumi: Conceptualization, Supervision, Writing - review & editing Minoru Inagaki: Conceptualization, Supervision, Writing - review & editing Kento Habe: Data curation, Formal analysis, Takuma Okabe: Data curation, Formal analysis Satoshi Tsuneda: Conceptualization, Supervision, Writing - review & editing. **Disclosure:** The authors declare no conflicts of interest associated with any financial or commercial arrangements of this manuscript.

## Abstract

Understanding the relationship between bacteriophage (phage) classification and target receptors is crucial for phage ecology and applied research. In this study, we compared 13 previously isolated *Escherichia coli* phages based on physiological characteristics, whole-genome sequences, and tail fiber protein phylogenetics. We improved our previously proposed physiological clustering method by optimizing the bacterial panel for host range assessment, implementing appropriate distance metrics for mixed data types, and applying silhouette coefficient analysis for objective determination of optimal cluster numbers. We combined genomic analysis and lipopolysaccharide (LPS) structural analysis of phage-resistant *E. coli* strains to identify target receptors of the phages. Complementation experiments further confirmed the direct involvement of identified genes in phage reception. The results revealed that phylogenetically distinct *E. coli* phages target different sites in the LPS R-core region (modified by WaaV, WaaW, WaaT, and WaaY), membrane proteins (NfrB, TolA, YhaH), or flagella. Our analysis revealed that, subtle chemical modifications of LPS (such as Heptose phosphorylation) were shown to be important for *E. coli* phage recognition. Furthermore, physiological characteristics, tail fiber phylogenetics, and whole genome analysis independently classified the phages with high correlation to target receptor specificity. The addition of three phages with known receptors further validated our approach. Our results suggest that grouping based on physiological characteristics (such as lysis dynamics and host range) and genotypes (tail fiber phylogenetics or whole genome analysis) independently classified phages with high correlation to target receptor specificity. Here, we elucidated the diversity and specificity of *E. coli* phage target receptors, providing new insights into the classification of *E. coli* phages and phage-host interactions.

**Importance:** Phage therapy is gaining attention as an alternative treatment for antibiotic-resistant bacteria. Developing effective phage cocktails requires combining phages with different target receptors, but traditional methods for identifying target receptors are labor-intensive. This study demonstrates that *E. coli* phages targeting the TK001 strain with different target receptors can be grouped based on their physiological characteristics, tail fiber sequences, or whole genomes. Our approach was enhanced through systematic bacterial panel selection for host range assessment, optimized distance metrics for physiological characteristics, and objective cluster determination using silhouette coefficient analysis for all three classifications. This insight can be used to more efficiently create diverse phage cocktails. Additionally, we identified phages targeting diverse sites, including different regions of LPS, membrane proteins, and flagella. These findings deepen our understanding of phage host recognition mechanisms, enabling the rapid preparation of effective phage cocktails and contributing to new advancements in bacterial infection treatment.

## Introduction

With the emergence of antibiotic-resistant bacteria, bacteriophage (phage) therapy-utilizing viruses that specifically infect bacteria - is gaining attention as an alternative treatment. Phages exhibit extremely high host specificity, which is primarily determined by their target receptors. For instance, phages that infect *Escherichia coli* are known to target membrane proteins, such as OmpC and LamB, or the lipopolysaccharide (LPS) (1). While bacteria can also become resistant to phages, using multiple phages that target different host receptors in a therapeutic cocktail has been shown to be effective at delaying and/or suppressing the emergence of phage-resistant bacteria (2)(3)(4)(5).

However, in the clinical setting, identifying the target receptors of each phage before creating a cocktail is impractical due to time constraints. For example, in a case of multidrug-resistant *Acinetobacter baumannii* infection reported by Schooley et al., the initial phage screening and selection was completed within 18 hours to address the patient’s critical condition, though additional phages were subsequently identified and administered to the patient. In such urgent cases, conventional methods based on receptor identification is impractical, and more rapid treatment decision-making is required (6). Traditionally, phage target receptor identification has involved a labor-intensive process of creating phage-resistant bacteria, conducting whole-genome comparative analysis, and identifying mutation sites (7)(8). While this methodology is reliable, it is unsuitable for clinical application due to its time and cost requirements. As such, in personalized phage therapy, phages are often combined in therapeutic cocktails without the knowledge of each phage’s receptor. If multiple phages target the same receptor, a single receptor mutation could render the cocktail ineffective.

In addition, while genomic approaches to phage characterization have become increasingly accessible for well-studied organisms like *E. coli*, receptor identification based on genomic data requires that the host bacterial genome is well-characterized, with comprehensive understanding of its surface structures and their genetic determinants. For emerging multidrug-resistant pathogens with less characterized genomes, such as *Stenotrophomonas maltophilia* or various non-model clinical isolates, the interpretation of genomic data for receptor identification remains challenging (9).

In our previous study, we proposed a clustering method based on physiological characteristics such as growth dynamics and host range. This method makes it possible to create effective phage cocktails without identifying the target receptors (10). However, at the time, we did not evaluate the validity of the proposed clustering methodology.

The present study aimed to validate and improve our previously proposed clustering method through comprehensive analysis of each phage’s target receptor. To address the limitations of our previous approach, we enhanced our methodology by implementing systematic bacterial panel selection for host range assessment, Gower distance calculation optimized for mixed data types, and silhouette coefficient analysis for objective cluster determination. Furthermore, we compared this improved physiological characteristic based classification method to other classification methods, including whole-genome sequencing and tail fiber protein phylogenetic analysis. Through these analyses, we aimed to evaluate how each classification method correlates with target receptors, with a particular focus on validating our refined physiological clustering approach. These findings are expected to contribute to rapid treatment decision-making in clinical settings and the standardization of phage therapy.

## Results

### Genomic characterization of phages

Whole genome sequencing and ANI analyses indicated that the phages represent three distinct taxonomic genera (Fig. S1, Table S1, S2, S3): ΦWec172, ΦWec174, and ΦWec177 belong to *Felixounavirus* (*Ounaviridae*); ΦWec189, ΦWec191, ΦWec193, and ΦWec196 are members of *Vequintaviridae* (*Vequintavirus*); and ΦWec179, ΦWec181, ΦWec186, ΦWec187, ΦWec188, and ΦWec190 are members of *Stephanstirmvirinae*. Among the *Stephanstirmvirinae* phages, ΦWec187 showed characteristics related to *Justusliebigvirus*, while our previous analysis indicated that ΦWec179, ΦWec181, ΦWec186, ΦWec188, and ΦWec190 represent a proposed novel genus “*Wecvirus*” (11).

### Whole genome analysis of TK001 wild type strain and mutation analysis of target receptor mutant strains

TK001 used in this study was isolated from the feces of a C57BL/6JJcl female mouse with dextran sodium sulfate induced colitis (MP Biomedicals, Soho, OH, USA) and was used as the host strain for phage isolation. Hybrid assembly of shotgun sequencing and long-read sequencing revealed that the complete genome of TK001 consists of 4,779,140 bp with a GC content of 50.79%, and its sequence type (ST) based on Multilocus sequence typing (MLST) analysis is 345.

Mutation analysis of 13 phage-resistant strains revealed frameshift mutations or stop codon insertions in genes encoding outer membrane/inner membrane proteins or LPS synthesis enzymes in each strain (Table 1). Among the 13 phage-resistant strains, 8 strains showed mutations in the genes of the LPS synthesis pathway, while the other 4 strains had mutations in genes related to membrane proteins or flagella. Notably, no target receptor mutants were obtained for ΦWec179. For ΦWec177, two resistant strains were analyzed and are distinguished as R177-1 and R177-2.

**Table 1.** Genes with frameshift or stop codon mutations in target receptor mutant *E. coli* strains.

| phage used to make strains resistant | strain | mutated gene(s) | description (role of genes) | mutation site in WT | mutation type | mutation site in WT amino acids (native length [AAs]) | the size of the expected gene product (WT size) [Da] |
| --- | --- | --- | --- | --- | --- | --- | --- |
| ΦWec172 | R172 | *waaV* | beta1,3-glucosyltransferase | 3934566 | Frame shift (A→.) | p.Y151Ifs*158 (327) | 17856.774 (38773.742) |
| ΦWec174 | R174 | *waaW* | UDP-galactose--(galactosyl) LPS alpha1,2-galactosyltransferase | 3935258 | Frame shift (.→T) | p.A290Dfs*295 (341) | 33723.99 (39312.395) |
| ΦWec177 | R177-1 | *waaV* | beta1,3-glucosyltransferase | 3934492 | Frame shift (A→.) | p.F175Lfs*181 (327) | 20786.28 (38773.742) |
| ΦWec177 | R177-2 | *waaV* | beta1,3-glucosyltransferase | 3934492 | Frame shift (A→.) | p.F175Lfs*181 (327) | 20786.28 (38773.742) |
|  |  | *wecA* | Undecaprenyl-phosphate alpha-N-acetylglucosaminyl 1-phosphate transferase | 4108976 | Frame shift (A→.) | p.A28Afs*28 (367) | 3647.418 (40938.761) |
| ΦWec181 | R181 | *manB-1* | Mannose-1-phosphate guanylyltransferase | 2178550-2178553 | Frame shift (CCTC→.) | p.A390_E391fs*432 (458) | 47602.579 (50839.022) |
|  |  | *waaT* | LPS 1,2-glucosyltransferase | 3937043 | Stop codon (C→T) | p.W269* (331) | 30982.807 (38497.428) |
| ΦWec186 | R186 | *flhD* | Flagellar transcriptional regulator FlhD | 2021475 | Ser→Pro (A→G) | p.S16P (156) | 13308.283 (13298.245) |
|  |  | *bcsG* | Cellulose biosynthesis protein BcsG | 3831107 | Stop codon (C→T) | p.Q542* (559) | 59871.85 (61986.166) |
| ΦWec187 | R187 | *bcsG* | Cellulose biosynthesis protein BcsG | 3831107 | Stop codon (C→T) | p.Q542* (559) | 59871.85 (61986.166) |
| ΦWec188 | R188 | *yhaH* | Inner membrane protein YhaH | 3405873-3405920 | Frame shift* | p.P66_A82del (121) | 12236.536 (14262.756) |
| ΦWec189 | R189 | *nfrB* | bacteriophage N4 adsorption protein B | 576511 | Stop codon (C→T) | p.W391* (745) | 45051.426 (85329.404) |
|  |  | *ompA* | Outer membrane protein A | 1046632-1046640 | Frame shift (CACCAAAAG→.) | p.A60_G63del (346) | 36906.975 (37182.278) |
| ΦWec190 | R190 | *waaG* | UDP-glucose:(heptosyl) LPS<br>alpha-1,3-glucosyltransferase | 3940586 | Stop codon (A→T) | p.L76*(374) | 8796.029<br>(42509.807) |
| ΦWec191 | R191 | *nfrB* | bacteriophage N4 adsorption<br>protein B | 576613 | Asp→Ala (T→G) | p.N357A (745) | 85285.395<br>(85329.404) |
|  |  | *tolA* | TolA | 767756 | Stop codon (C→T) | p.Q269* (422) | 7295.244<br>(43138.074) |
| ΦWec193 | R193 | *waaY* | LPS core heptose (II) kinase<br>RfaY | 3936259 | Stop codon (C→A) | p.E196* (230) | 22793.077<br>(26975.989) |
| ΦWec196 | R196 | *waaY* | LPS core heptose (II) kinase<br>RfaY | 3936259 | Stop codon (C→A) | p.E196* (230) | 22793.077<br>(26975.989) |

### Identification and validation of phage receptor genes through efficiency of plating (EoP) of phages against target receptor mutant strains and Keio collection, complementation, and structural prediction

The EoP of each phage against TK001 and target receptor mutant strains was evaluated (Fig. 1). A decrease in EoP to less than 10-fold (log(EoP) < −1) was defined as a reduction in infection efficiency. All phages could infect *bcsG* mutant (R187), a mutation common to R186 (*fihD* and *bcsG* mutant); therefore, *bcsG* was not considered to be involved in target recognition.

**Fig. 1.**
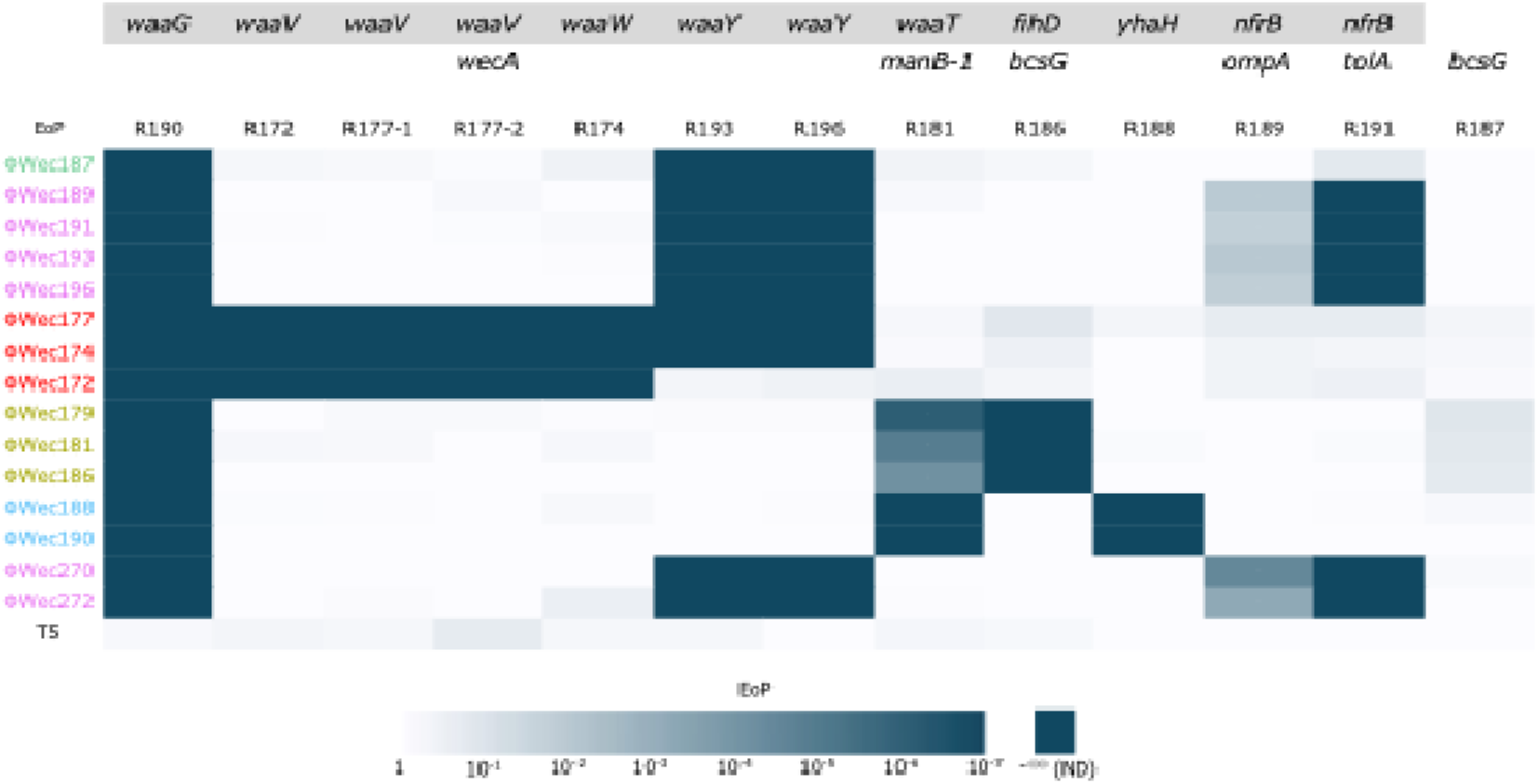
Heatmap of infectivity of each phage against phage-resistant *E. coli* strains

Regarding LPS synthesis genes, none of the phages could infect the *waaG* mutant (R190). ΦWec172, 174, and 177 lost their ability to infect *waaW* mutant (R174) and *waaV* mutants (R172, R177-1, and R177-2). ΦWec174, 177, 187, 189, 191, 193, and 196 could not infect *waaY* mutants (R193 and R196). ΦWec179, 181, 186, 188, and 190 lost infectivity against *waaT* and *manB-1* double mutant (R181).

For membrane protein and flagella genes, ΦWec179, 181, and 186 could not infect *fihD* mutant (R186). ΦWec188 and 190 lost infectivity against *yhaH* mutant (R188). ΦWec189, 191, 193, and 196 could not infect *nfrB* and *ompA* double mutant (R189) and *nfrB* and *tolA* double mutant (R191).

Using ΦWec187, 189, 191, 193, and 196, which can infect the K-12-derived BW25113 strain (wild-type of the Keio collection), we investigated their ability to infect Keio collection strains with single gene deletions corresponding to the mutations found in phage-resistant strains (Fig S2). ΦWec187, 189, 191, 193, and 196 all failed to infect Δ*nfrB*, Δ*waaG*, and Δ*waaF* strains. Additionally, ΦWec189, 191, 193, and 196 could not infect the Δ*nfrA* strain, which is genetically redundant with *nfrB*. Based on these results, the genes necessary for infection by each phage were classified into six types (Table 2). Types a and b were very similar, differing only in the presence or absence of a *waaY* mutation.

**Table 2.**
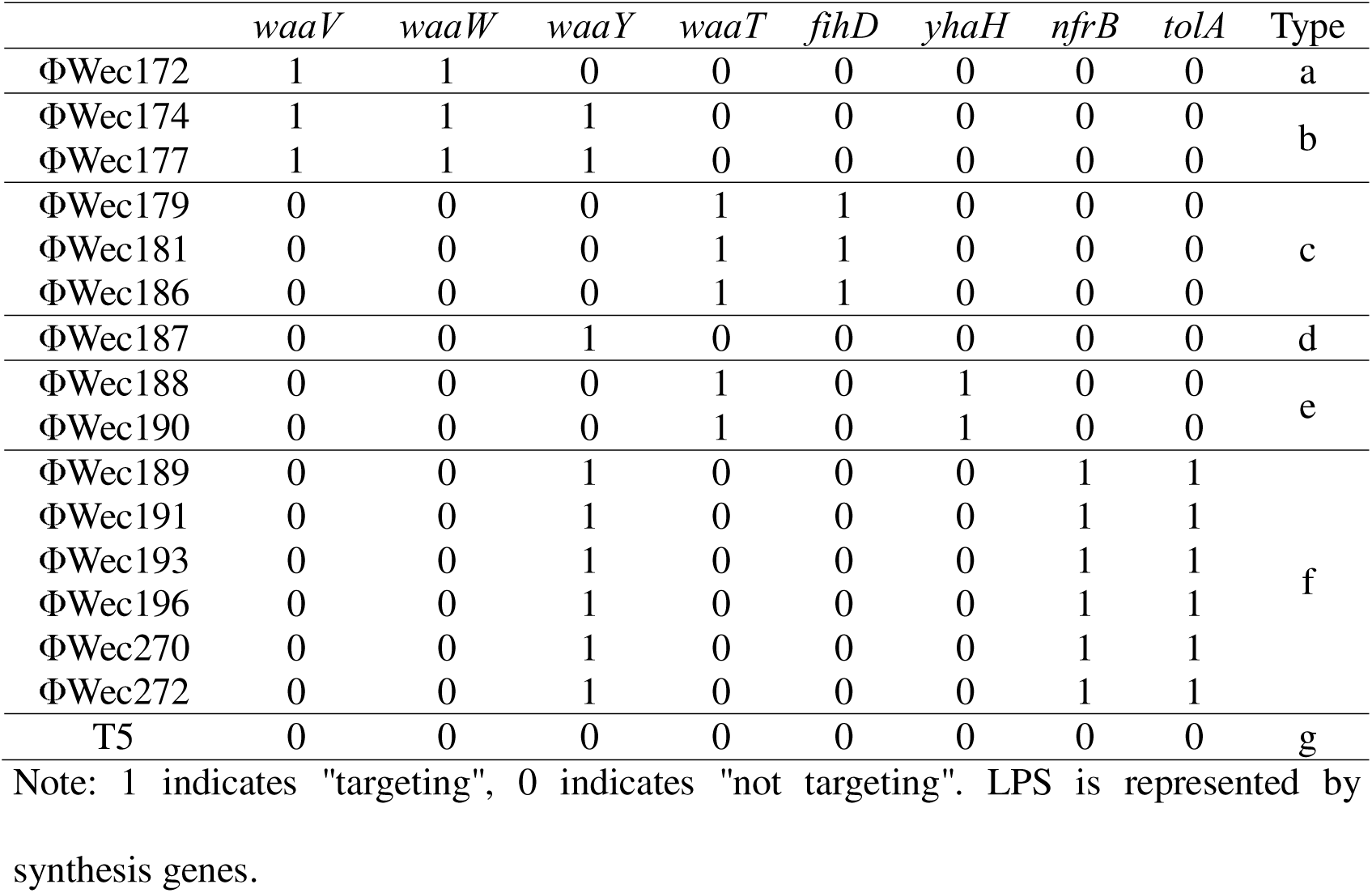
Genes affecting target cell recognition by each phage.

To further validate our receptor identification, we performed two additional analyses. First, we tested three additional phages (ΦWec270, ΦWec272, and T5) against our panel of receptor mutant strains. ΦWec270 and ΦWec272, which belong to the same genus as ΦWec189, 191, 193, and 196 based on previous genomic analyses, showed identical infection patterns to these phages, confirming they share the same target receptor requirements including both NfrB and phosphorylated heptose in LPS (mediated by WaaY). In contrast, T5, which is known to target FhuA in *E. coli* K-12 (12)(13) and is taxonomically distinct from all other phages used in this study, remained infective against all our receptor mutant strains, indicating it recognizes a different receptor not affected by our mutations.

Second, we conducted complementation experiments by expressing wild-type genes in the corresponding mutant strains using plasmid vectors. When the wild-type versions of the mutated genes were reintroduced, infection sensitivity was restored in most cases, confirming the direct involvement of these genes in phage reception (Fig. 2). Specifically, complementation of *waaW*, *waaG*, *waaT*, *waaY*, *nfrB*, *flhD*, and *yhaH* mutations restored sensitivity to the respective phages that lost infectivity against these mutants.

**Fig. 2.**
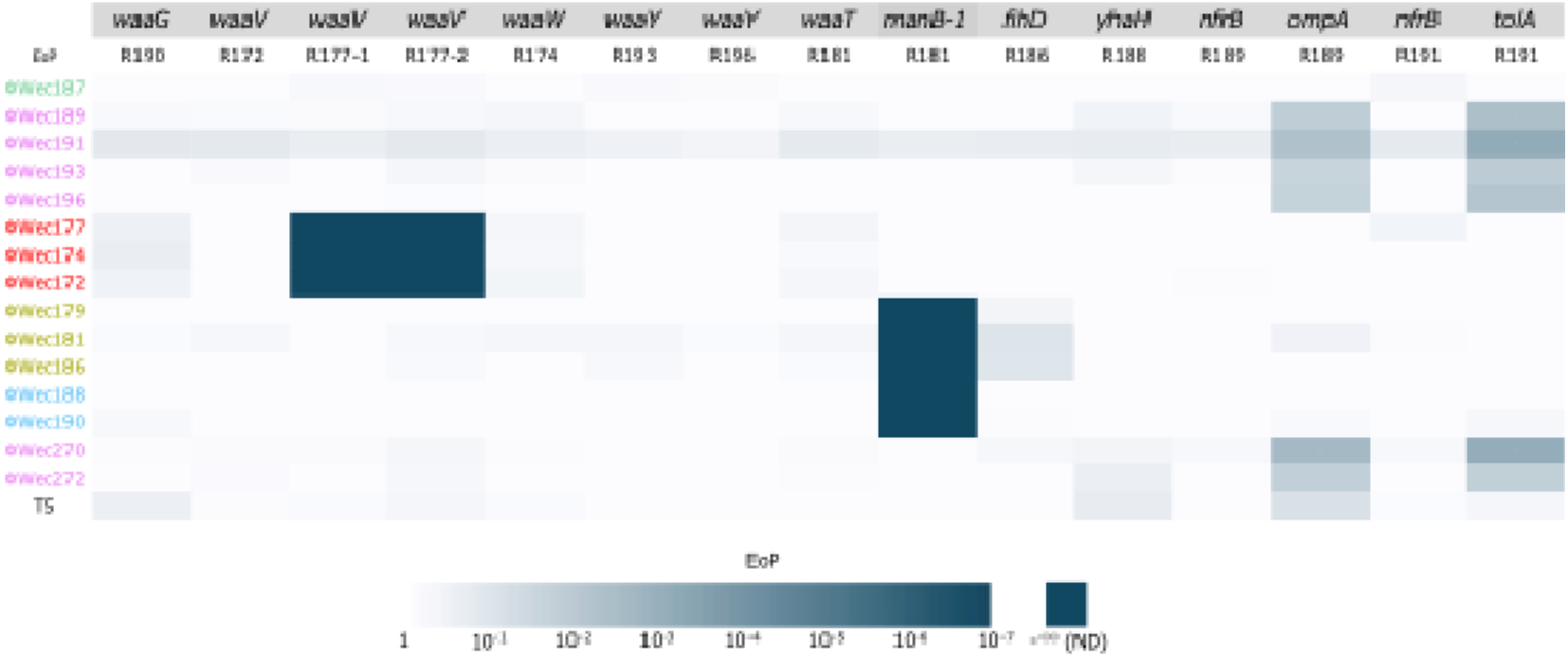
Heatmap of infectivity of each phage against complemented resistant mutants

Interestingly, complementation results for *waaV* mutations were variable. Introduction of wild-type *waaV* into the R172 mutant (with deletion at position 451 bp) restored phage sensitivity, while complementation failed to restore sensitivity in R177-1 and R177-2 mutants (both with deletion at position 525 bp). To investigate whether this was due to the mutant protein interfering with wild-type function or due to polar effects on downstream genes, we expressed the mutant *waaV* variants in wild-type TK001. Neither variant affected EoP, suggesting that the different complementation outcomes might be related to secondary effects specific to the R177-1/2 genetic background.

Structural predictions of LPS synthesis enzymes using AlphaFold2 and AlphaFold3 were used to visualize the molecular impacts of the identified mutations (detailed analyses in Supplementary Results and Discussion, Fig. S3-9). For structural analysis, we referenced experimentally determined crystal structures when available (e.g., *E. coli* K-12 WaaG, PDB 2IW1) to analyze ligand interactions and validate our AlphaFold predictions. The analysis revealed that mutations in *waaG*, *waaV*, *wecA*, and *manB-1* resulted in extensive truncations (45-92% of the protein). Mutations in *waaW*, *waaT*, and *waaY* resulted in more limited truncations (14-19% of the protein), primarily affecting C-terminal regions rich in basic amino acids while preserving most catalytic residues.

Host range specificity of each phage was evaluated against target receptor mutant strains using spot tests. The heatmap displays EoP values calculated relative to the wild-type TK001 strain. White indicates no reduction in infectivity (EoP = 1), and progressively darker cyan shades represent decreasing infectivity levels, with dark cyan signifying complete loss of infectivity (not detected, ND). Each mutant strain is labeled with the specific gene(s) containing frameshift or stop codon mutations, with gray backgrounds indicating LPS biosynthesis genes. Phages are color-coded by classification groups and arranged to highlight infection pattern similarities within groups. For ΦWec179, 181, and 186 against R181, 4 out of 7 trials were ND (Not Detected), and the log (EoP) value in the table is the average of the remaining 3 trials.

Infectivity recovery was evaluated by introducing wild-type genes into the corresponding resistant mutant strains. The heatmap displays EoP values calculated relative to the wild-type TK001 strain carrying empty vector control. White indicates no reduction in infectivity (EoP = 1), and progressively darker cyan shades represent decreasing infectivity levels, with dark cyan signifying complete loss of infectivity (not detected, ND). Gray backgrounds indicate LPS biosynthesis genes. The order and color coding of phages is maintained consistent with Fig. 1 for ease of comparison. Phages of the same color show similar infectivity patterns against both the resistant mutants and their complemented derivatives.

### LPS structure of TK001 wild type and phage resistant strains

Mutations in genes involved in the LPS synthesis pathway were identified in many phage-resistant *E. coli* strains. Most of the LPS-related enzymes mutated in the target receptor mutant strains were associated with R-core synthesis, while *wecA* and *manB-1* were genes for synthesizing sugars that constitute the O-antigen. Each *E. coli* strain possesses one of five R-core types, and the analysis suggested that the R-core of TK001 was the R1 type (Fig. 3a, Fig. S10a) (14). Variations in the R-core structure were suggested to arise from deficiencies in the R-core synthesis pathway genes (Fig. S10b). The KEGG mapper results indicated the presence of both *waaR* and *waaT* genes, as these genes share the same KEGG Orthology (KO) identifier in the database, though they encode functionally distinct enzymes. However, examination of the actual genome annotation revealed that TK001 possesses only the *waaT* gene and not *waaR*, suggesting that the cascade beyond *waaR* is not active.

**Fig. 3.**
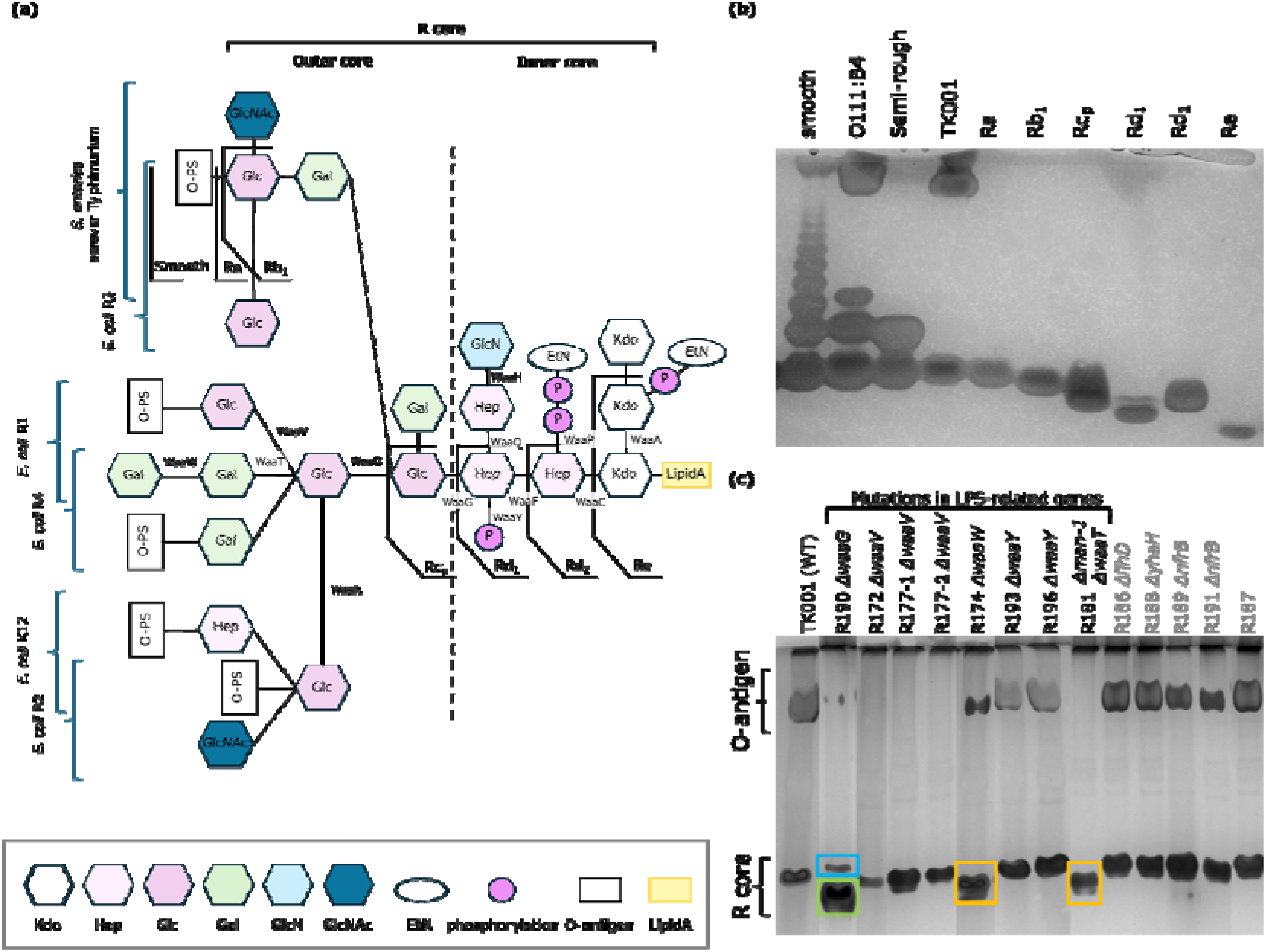
LPS structural analysis and comparison across reference strains and TK001 mutants (a) Schematic representations of LPS structures of *Salmonella enterica* serovar Typhimurium and *E. coli* R1, R2, R3, R4 and K12. The predicted LPS structure of TK001 and associated biosynthesis enzymes is also shown. (b) DOC-PAGE analysis comparing LPS from TK001 with reference strains. “Smooth” is LPS from *S. enterica* serovar Typhimurium, showing ladder-like bands characteristic of smooth strains with varying numbers of repeats of O-antigen. “O111:B4” is LPS from *Escherichia coli* O111:B4, showing a band pattern characteristic of smooth strains with relatively compact long repeat units. “Semi-rough” is LPS from *S. enterica* serovar Typhimurium SL1034 (semi-rough chemotype). “Ra” is Ra chemotype. Same as others. (c) DOC-PAGE analysis of LPS from TK001 phage-resistant mutant strains. The order has been rearranged for ease of viewing. Orange and green highlights show the downward shift of the lower bands compared to the wild type. Blue highlights indicate the presence of an additional band distinct from the band marked by the green highlight.

DOC-PAGE analysis of LPS structure revealed that TK001 is similar to *E. coli* O111:B4, suggesting the presence of LPS with a relatively uniform, long O-antigen repeat unit in addition to the R-core (Fig. 3b). Strains with mutations in genes unrelated to LPS synthesis showed the same band pattern in DOC-PAGE as TK001, indicating no change in LPS structure (Fig. 3c, Fig. S11). In *waaV* mutants (R172, R177-1, and R177-2), O-antigen (the upper band in DOC-PAGE) completely disappeared. The DOC-PAGE profile of the *waaW* mutant (R174) showed changes in the shape of the upper band and a downward shift in the R-core band position. The *waaT* mutant (R181) showed complete disappearance of the upper band and a downward shift in the R-core band position. The *waaG* mutant (R190) showed near disappearance of the upper band and separation of the R-core band into two. No clear changes in band pattern were observed in *waaY* mutants (R193 and R196).

### Improved methodology for physiological clustering of phages

To enhance the rigor of our previously proposed classifying method based on physiological characteristics, we implemented several methodological improvements. First, we optimized the bacterial panel for host range assessment by analyzing an infection matrix of 58 phages, which were primarily those used in our previous research (15), against 41 *Enterobacteriaceae* strains (Table S4, S5). Using a greedy algorithm, we selected strains that collectively maximized the number of distinct infection patterns observed. This approach identified 16 strains (Table 3) that collectively revealed 39 distinct infection patterns, providing a more systematic basis for host range evaluation than previously used.

**Table 3.** Bacterial panel selected for optimized host range assessment Comparison of target receptors with various classification methods.

| Rank | *Enterobacteriaceae* strain | Additional discriminatory patterns | Cumulative discriminatory patterns |
| --- | --- | --- | --- |
| 1 | TK001 | 2 | 2 |
| 2 | MG1655 | 2 | 4 |
| 3 | BL21 | 3 | 7 |
| 4 | ESBL953 | 5 | 12 |
| 5 | NBRC102203 | 6 | 18 |
| 6 | ESBL983 | 4 | 22 |
| 7 | ATCC43888 | 3 | 25 |
| 8 | TOP10F' | 2 | 27 |
| 9 | ESBL946 | 2 | 29 |
| 10 | ESBL955 | 2 | 31 |
| 11 | ESBL1054 | 2 | 33 |
| 12 | *E. fergusonii* NBRC102419 | 2 | 35 |
| 13 | SUTL-3 | 1 | 36 |
| 14 | ESBL920 | 1 | 37 |
| 15 | ESBL1013 | 1 | 38 |
| 16 | *S. Pullorum* NBRC3163 | 1 | 39 |

Note: All of them are *E. coli* other than *E. fergusonii* and *S. Pullorum*

For clustering analyses, we implemented Gower distance instead of Euclidean distance to appropriately handle both qualitative infection data and quantitative physiological measurements. To objectively determine the optimal number of clusters, we applied silhouette coefficient analysis, which identified k=4 as the optimal number of clusters for our original set of 13 phages (Fig. 4a).

**Fig. 4.**
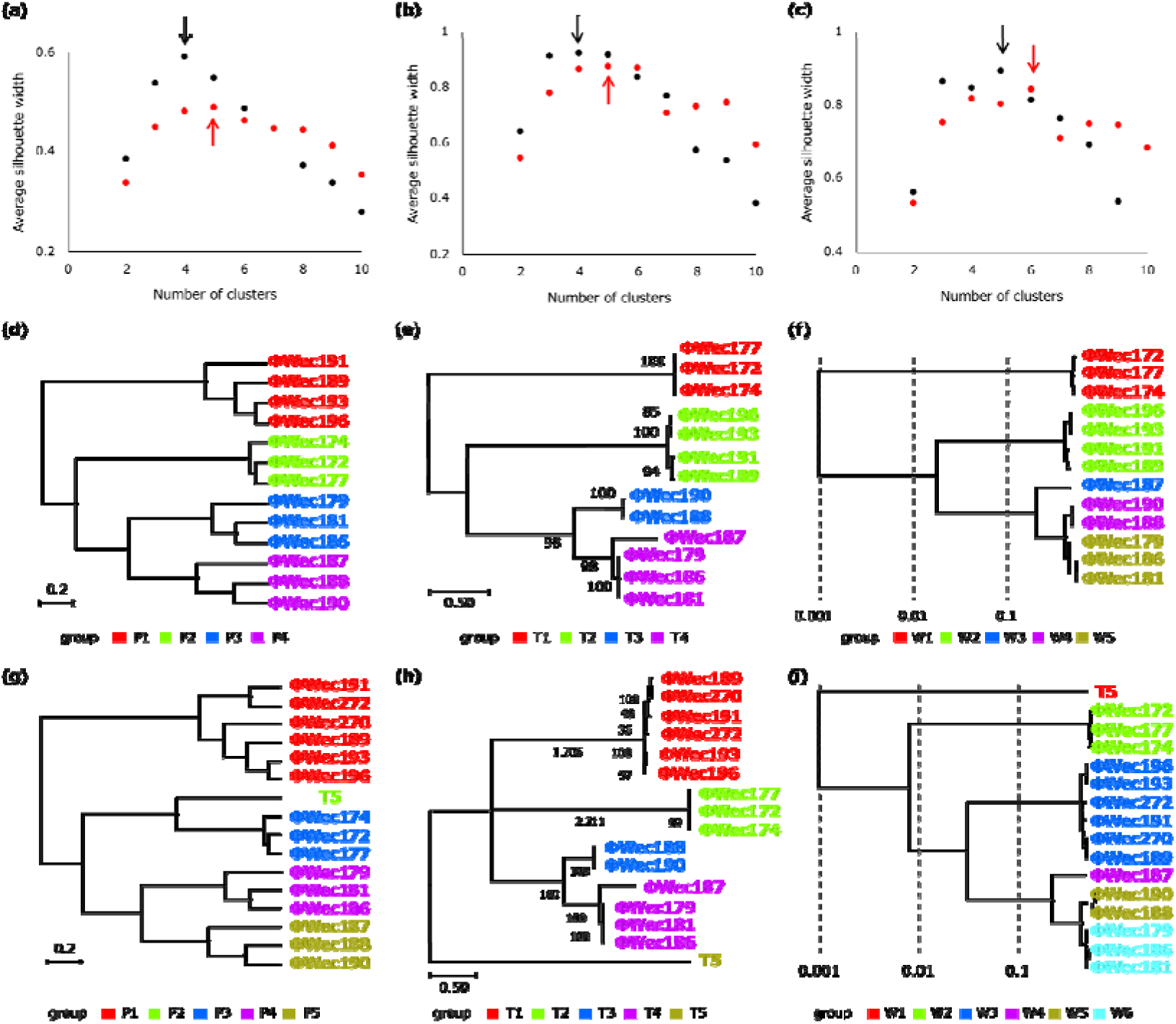
Silhouette coefficient analysis and hierarchical clustering of phages by different methods (a-c) Silhouette coefficient analysis for determining optimal cluster numbers based on (a) physiological characteristics, (b) tail fiber phylogeny, and (c) whole genome phylogeny. The plots show average silhouette width values for different numbers of clusters (k) from 2 to 10. Black plots represent analysis of 13 original phages, while red plots show results after addition of three supplementary phages (ΦWec270, ΦWec272, and T5). Arrows indicate the maximum silhouette coefficient values, representing the optimal number of clusters for each method. Note that in panel (c), results for k=10 are not shown as the coefficient value diverged to negative infinity for whole genome phylogeny. (d-f) Hierarchical clustering dendrograms for the original 13 phages based on (d) physiological characteristics, (e) tail fiber phylogeny, and (f) whole genome phylogeny. (g-i) Hierarchical clustering dendrograms after addition of three supplementary phages based on (g) physiological characteristics, (h) tail fiber phylogeny, and (i) whole genome phylogeny. Colors in dendrograms (d-i) represent clusters determined using the optimal number identified by silhouette coefficient analysis in panels (a-c).

The resulting clustering (Fig. 4d) revealed four distinct phage groups: P1 (ΦWec189, 191, 193, 196), P2 (ΦWec172, 174, 177), P3 (ΦWec179, 181, 186), and P4 (ΦWec187, 188, 190). When additional phages with known receptors (ΦWec270, ΦWec272, and T5) were included, silhouette analysis identified k=5 as the optimal number, with T5 forming its own distinct group (P5), while ΦWec270 and ΦWec272 clustered with group P1 (Fig 4a, g).

To evaluate how our improved physiological clustering correlates with target receptor specificity, we compared groupings based on physiological characteristics, tail fiber phylogeny, and whole genome phylogeny with grouping based on target receptors. Using silhouette coefficient analysis, we determined the optimal cluster numbers for tail fiber phylogeny (k=5) and whole genome phylogeny (k=4) for the original 13 phages (Fig. 4a-c).

The grouping results from the three classification methods are summarized in Fig. 4d-f. While each method produced somewhat different groupings, there were notable consistencies. For example, ΦWec172, ΦWec174, and ΦWec177 were consistently grouped together across all three methods (P2, T1, W1). Similarly, ΦWec189, ΦWec191, ΦWec193, and ΦWec196 formed a consistent group (P1, T2, W2).

When including the three additional phages, the optimal cluster numbers increased to 5, 6, and 5 for physiological, tail fiber, and whole genome classifications, respectively.

The similarity between these groupings and the phage groups based on target receptors was evaluated using the Adjusted Rand Index (ARI), which measures the agreement between two clustering methods (Table 4). All three classification methods showed high correlation with the grouping based on target receptors, both before and after the addition of the three new phages. Notably, the addition of phages with distinct receptor specificity improved the ARI values for all three methods, demonstrating the robustness of these classification approaches.

**Table 4.** ARI Values of Similarity between each Clustering Method with Target Receptor-Based Grouping Before and After Additional Phages.

| Comparison Pair | ARI |  |
| --- | --- | --- |
|  | 13 phages | 16 phages |
| Physiological vs Target receptor | 0.82 | 0.89 |
| Tail fiber vs Target receptor | 0.78 | 0.86 |
| Whole genome vs Target receptor | 0.90 | 0.94 |

The whole genome-based classification consistently showed the highest similarity with target receptor grouping (ARI = 0.90 for 13 phages, increasing to 0.94 for 16 phages). Physiological characteristic-based clustering showed substantial improvement with the addition of new phages (ARI increasing from 0.82 to 0.89), demonstrating its ability to accurately classify phages with diverse receptor specificities.

Furthermore, we evaluated the similarity between the different classification methods themselves (Table 5). All three methods showed high concordance with each other, with ARI values ranging from 0.80 to 0.95. The addition of the three phages further increased the agreement between methods, indicating that as the diversity of phages increases, these different classification approaches tend to converge in their grouping patterns.

**Table 5.** ARI Values of Similarity between Three Different Classification Methods Before and After Additional Phages.

| Comparison Pair | ARI |  |
| --- | --- | --- |
|  | 13 phages | 16 phages |
| Physiological vs Tail fiber | 0.80 | 0.87 |
| Physiological vs Whole genome | 0.91 | 0.95 |
| Tail fiber vs Whole genome | 0.87 | 0.92 |

These results demonstrate that all three classification methods reflect the underlying biological relationships between phages, particularly their target receptor specificity. Importantly, the physiological characteristic-based classification, which can be determined without extensive genomic analysis, provides a reliable basis for estimating both genetic relationships and receptor specificity of phages across a diverse range of phage types.

## Discussion

While phage therapy using a cocktail of phages targeting different receptors is known to be effective, methods to identify phage target receptors remain limited and time-consuming. In our previous study, we demonstrated that effective phage cocktails could be prepared by grouping phages based on their physiological characteristics, but the actual receptors of the phages were not confirmed (9). In this study, we validated our previous clustering method through detailed analysis of each phage’s target receptor. Our findings demonstrate that phages with different target receptors can be grouped based on their physiological characteristics, tail fiber sequences, or whole genomes. These results provide a foundation for more efficient creation of diverse phage cocktails and deepen our understanding of phage-host interactions, enabling rapid preparation of effective therapeutic combinations.

### Phage target receptors

In this study, we identified the target receptors of each phage in detail by combining the analysis of phage-resistant strains with LPS structure analysis. These results reveal the diversity and specificity of phage host recognition mechanisms and provide new insights into the relationship between LPS structure and phage infectivity (Fig. 5).

**Fig. 5.**
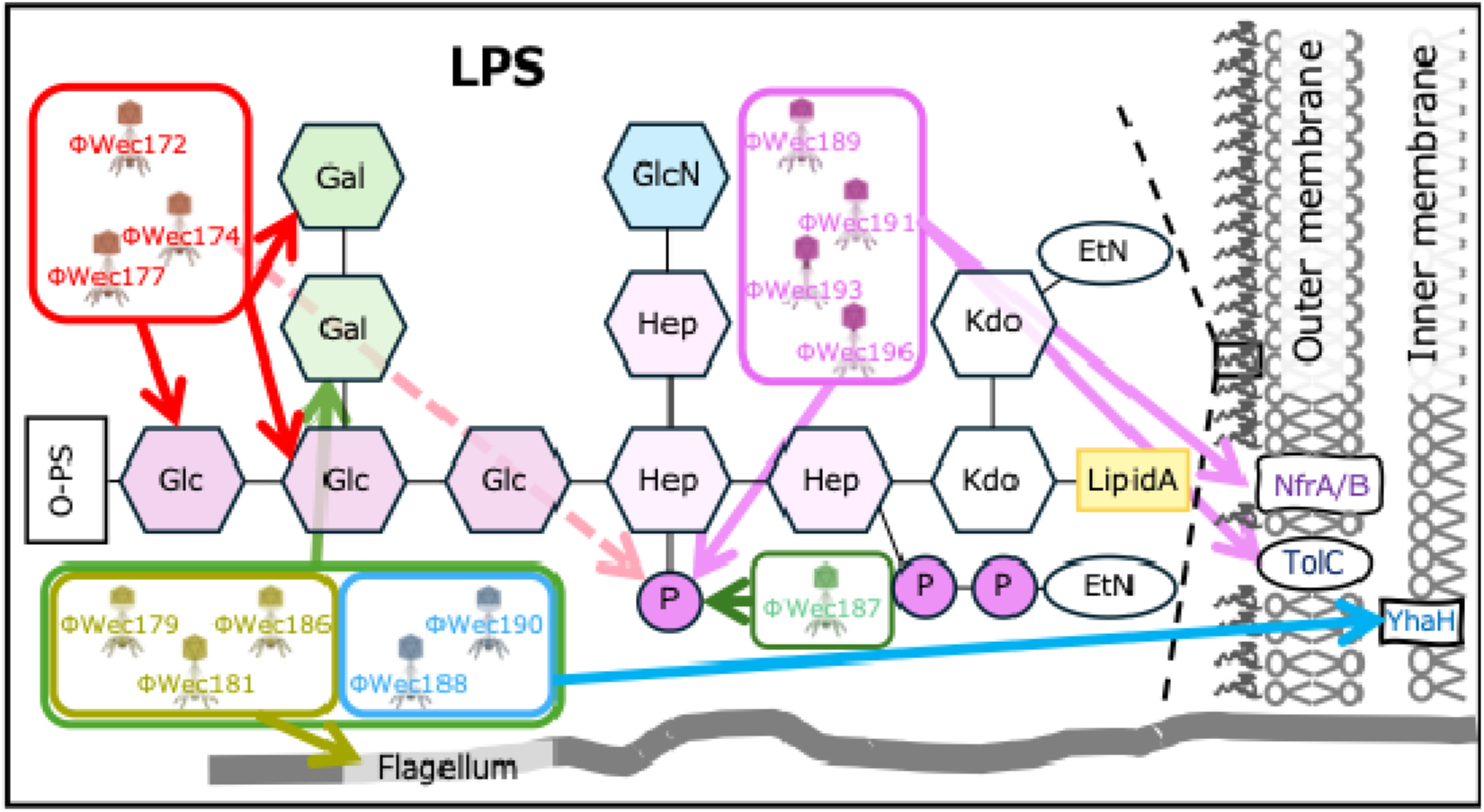
Target receptors of each phage

The results of LPS structure analysis clearly show the specific effects of each gene mutation on LPS structure. The complete disappearance of the upper band in DOC-PAGE for *waaV* mutants (R172, R177-1, and R177-2) strongly suggests that in the TK001 strain, the O-antigen is attached to the Glcβ added by WaaV. Additionally, the downward shift of the R-core bands in *waaW* and *waaT* mutants (R174 and R181) indicates that the deficiency of these genes results in a shorter R-core length. Interestingly, in the *waaG* mutant (R190), the upper band almost disappeared, and the R-core band separated into two. Since *waaG* is an early gene in outer core synthesis, this result suggests that while R-core synthesis was incomplete, some compensatory mechanism may have led to the expression of a unique LPS, consistent with previous observations by Klein et al. who demonstrated that *waaG* deletion in *E. coli* K-12 resulted in increased incorporation of a third Kdo residue and rhamnose, leading to accumulation of different structures of glycoforms (16). On the other hand, no clear changes in band pattern were observed in *waaY* mutants (R193 and R196), indicating that WaaY, which is involved only in Hep phosphorylation, has little effect on the overall length of LPS but creates functionally critical modifications for specific phage recognition. Notably, although ΦWec172, 174, and 177 were suggested to be similar in our previous report (10) due to their common isolation source, similar lysis range, and over 99% homology in ANI-based similarity evaluation (Table S4), ΦWec172 did not require WaaY’s function, unlike ΦWec174 and 177. These phages have two types of tail fibers: Tail fiber 1 consisting of 844 amino acid residues and Tail fiber 2 consisting of 390 amino acid residues. There are the N777S mutation in Tail fiber 1 of ΦWec174 and the K286E mutation in Tail fiber 2 of ΦWec177. These differences suggest that ΦWec172 may interact with LPS differently from the other two phages and may not require Heptose phosphorylation by WaaY. This observation demonstrates the complexity and flexibility of phage receptor recognition and may provide an example of adaptation strategies in phage-host interactions. However, as no unique mutations were found in these tail fibers for ΦWec172 alone, further genomic analysis, structural analysis, or mutation experiments are necessary to elucidate the mechanism.

The changes in LPS structure were closely related to the infectivity of each phage. The most important finding was that all phages were unable to infect the *waaG* mutant (R190), clearly demonstrating that LPS is essential for infection by all phages used in this study.

Our structural prediction analyses revealed how different types of mutations in LPS biosynthesis genes lead to diverse effects on receptor function. For extensive truncations in WaaG, WaaV, WecA and ManB-1, the complete loss of catalytic domains explains the absence or severe disruption of LPS structures observed in DOC-PAGE. In contrast, for WaaW, WaaT, and WaaY mutations, the truncations appeared less extensive but potentially significant to function. These mutations preserved most catalytic residues but affected C-terminal regions containing clusters of basic amino acids. These positively charged regions likely interact with the negatively charged LPS, suggesting how even limited truncations could substantially alter substrate binding capabilities without complete loss of enzymatic activity.

The WaaY mutations provided particularly compelling evidence for this mechanism. DOC-PAGE analysis showed no visible changes in overall LPS length, suggesting the core structure remained intact. However, phage infection experiments clearly demonstrated functional changes, as certain phages (ΦWec189, ΦWec191, ΦWec193, and ΦWec196) could not infect these mutants. Complementation with wild-type WaaY restored phage sensitivity, confirming the direct relationship between the mutation and phenotype. Our structural predictions explain this apparent discrepancy - WaaY’s kinase activity phosphorylates heptose residues without altering core length, creating negatively charged regions that serve as critical recognition sites for specific phages. This demonstrates the remarkable discriminatory capacity of phages, which can evolve to recognize highly specific chemical modifications on the bacterial surface.

Furthermore, interesting results were obtained regarding the target receptor recognition of ΦWec172, 174, and 177. These phages could not infect the *waaW* mutant but could infect the strain with a mutation in WaaT, which functions just before WaaW in the LPS synthesis pathway. In the LPS synthesis pathway, WaaT transfers Galactose, followed by WaaW transferring second Galactose. This result suggests that these phages primarily target the Galactose transferred by WaaW. However, the fact that they can also infect the WaaT mutant suggests that similar structures, such as Glucose transferred by WaaO, may serve as alternative targets. This flexible recognition mechanism might be one of the phages’ strategies for adapting to their hosts.

Our variable results in complementing *waaV* mutations highlight the complexity of LPS biosynthesis pathways. The successful complementation in R172 (deletion at position 451 bp) but not in R177-1 and R177-2 (deletions at position 525 bp) suggests that the position of the mutation within *waaV* may lead to different effects on downstream genes or LPS assembly mechanisms. The failure of mutant *waaV* variants to affect EoP when expressed in wild-type TK001 indicates that these mutations are likely recessive and do not interfere with wild-type function, pointing to potential secondary effects in the genetic background of specific mutants.

The addition of phages with known receptors (ΦWec270, ΦWec272, and T5) further validated our approach. ΦWec270 and ΦWec272, which share the same genus as ΦWec189, 191, 193, and 196, showed identical infection patterns, confirming they target the same receptor structures requiring both NfrB and phosphorylated heptose in LPS. In contrast, T5, which targets FhuA in *E. coli* K-12, remained infective against all our receptor mutant strains, indicating it recognizes a receptor not affected by our mutations. This diversity in receptor recognition mechanisms demonstrates the importance of using multiple phages targeting different host receptors in therapeutic applications.

The receptor diagram illustrates the diversity of receptors including LPS components, membrane proteins, and flagella. NfrA/B are depicted together as they represent functionally linked membrane proteins that cooperatively serve as phage receptors (18). Arrows extending from each phage indicate their target receptors. The dotted red arrow from ΦWec172 indicates that this phage does not require WaaY-mediated heptose phosphorylation for infection, despite being grouped with ΦWec174 and ΦWec177 based on genomic and physiological characteristics. This suggests subtle differences in receptor recognition mechanisms even among closely related phages.

### Phage classification methods

In this study, we compared three approaches for phage classification: physiological characteristics, tail fiber phylogeny, and whole genome phylogeny. We enhanced our previously proposed physiological classification method through several methodological improvements, including optimized bacterial panel selection, implementation of Gower distance for mixed data types to properly handle both qualitative infection data and quantitative measurements. Additionally, we applied silhouette coefficient analysis for objective determination of optimal cluster numbers across all classification methods, eliminating the subjective nature of cluster number selection that was a limitation in previous work.

Comparison of these methods with grouping based on target receptors showed that all methods performed better than random classification, with Adjusted Rand Index (ARI) values well above zero. All methods showed high agreement with target receptor-based classification, demonstrating strong interchangeability between these approaches. The improvement in ARI values after adding phages with distinct receptor specificities (ΦWec270, ΦWec272, and T5) further validates the robustness of our classification approaches. The proper clustering of T5 as a separate group across all methods demonstrates their ability to distinguish phages with unique receptor specificities, while the consistent grouping of ΦWec270 and ΦWec272 with phages sharing the same receptors confirms the methods’ sensitivity to receptor-related characteristics.

While overall agreement between methods was high, some phages showed variable classification depending on the method used. Specifically, ΦWec187 was grouped differently across methods - clustering with ΦWec188 and ΦWec190 in physiological-based grouping, with ΦWec179, 181, and 186 in tail fiber analysis, and forming its own distinct group in whole genome classification. This variability persisted even after adding new phages to the analysis, suggesting that ΦWec187 possesses a unique combination of characteristics that are captured differently by each method. Such observations highlight that each classification approach may emphasize different aspects of phage biology.

Importantly, none of the methods completely matched the classification based on target receptors. This suggests that phage target receptor specificity is a complex characteristic that cannot be fully explained by a single classification criterion. The recognition of target receptors is likely determined by not only specific regions of phage tail fibers but also other structural proteins. Furthermore, subtle differences such as structural elements may play significant roles that cannot be fully captured by whole genome analysis or physiological characteristics alone.

For example, studies on bacteriophage T4 have revealed that structures such as the collar and whiskers act as environment-sensing devices, regulating the retraction of long tail fibers under unfavorable conditions and thus preventing infection, demonstrating that phage-host recognition can involve structural elements beyond tail fibers to modulate infection processes (17). These results highlight the challenges in phage classification and functional prediction, and reflects the complexity of phage-host coevolution.

Nevertheless, this study demonstrated that our improved clustering based on physiological characteristics is an effective method that reflects phage target receptor specificity to a high degree. The high correlation between classifications based on phenotypic data and genomic information supports the validity of our approach for practical applications in phage selection, providing an important foundation for the development of effective phage therapy.

### Limitations and future directions

While our findings provide strong support for the validity of physiological characteristic-based clustering as a proxy for target receptor diversity, our analyses were conducted using a single *E. coli* strain (TK001) as the primary host, and the identified receptor mutations may be specific to this genetic background. The high agreement between our different classification methods suggests that physiological characteristics can serve as reliable indicators of genetic relationships and receptor specificity. This has important practical implications, particularly for phages targeting bacterial species with limited genomic characterization. As noted in our introduction, while genomic approaches have become increasingly accessible for well-studied organisms like *E. coli*, receptor identification based on genomic data requires well-characterized bacterial genomes. For emerging multidrug-resistant pathogens with less characterized genomes, our physiological classification approach could provide a practical alternative that does not depend on extensive genomic information.

In conclusion, this study provides important insights into *E. coli* phage target receptors and the relationship between various classification methods. Our findings reveal that phage host recognition mechanisms are diverse and specific, with many phages recognizing specific structures of LPS, while others target membrane proteins or flagella. Through robust molecular analysis of 13 phages, including detailed characterization of their physiological features, whole genome sequences, and target receptors, we have definitively validated our previously proposed classification method by demonstrating its correlation with actual phage target receptors.

While our findings are based on *E. coli* phages, this approach has potential broader applications. Further studies with phages infecting other bacterial species will be necessary to confirm the universal applicability of these methods. Nevertheless, in clinical settings where conventional receptor identification is impractical and rapid decision-making is crucial, our validated classification approach offers powerful tools for creating effective phage cocktails that can suppress the emergence of phage-resistant bacteria, marking a significant advancement in the field of phage therapy.

## Materials and methods

### Reagents and bacteria

Luria–Bertani (LB) medium (BD Difco, Franklin Lakes, NJ, USA) and the soft agar containing 0.5% agar were used for bacterial and phage cultivation. *E. coli* strain TK001, originally isolated from dextran sodium sulfate (MP Biomedicals, Soho, OH, USA)-induced colitis mouse feces, was used as the host for phage screening in this study (10).

### Phage preparation

Thirteen phages (ΦWec172, ΦWec174, ΦWec177, ΦWec179, ΦWec181, ΦWec186, ΦWec187, ΦWec188, ΦWec190, ΦWec189, ΦWec191, ΦWec193, and ΦWec196) were propagated using *E. coli* TK001 as the host through the plate lysate method.

For validation experiments, three additional phages were included. ΦWec270 and ΦWec272, previously characterized in our taxonomic study (15), and bacteriophage T5 (NBRC 20005) from the Biological Resource Center, National Institute of Technology and Evaluation (NBRC), were used. These three phages can infect TK001, but were propagated using *E. coli* MG1655 (CGSC #6300) as the host using the same plate lysate method.

Briefly, 100 μL of overnight-cultured TK001 and 100 μL of phage suspension (10^4^ plaque-forming unit (PFU)/mL) were added to soft agar (0.5% agar) and overlaid on agar plates. After overnight incubation at 37 °C, 5 mL of SM buffer (0.1 M NaCl, 8 mM MgSO_4_•7H_2_O, 50 mM Tris-HCl buffer [pH=7.5], 0.1% gelatin) was added to each plate and subjected to orbital shaking (approximately 250 rpm, 2-3 hours). The supernatant was centrifuged (9,000 rpm, 15 min, 4 °C) to separate solid and liquid components. The supernatant was transferred to a new tube, and 1 mL of chloroform was added. After vortexing and settling, and purified phage was decanted and served as the phage stock.

### Generation and selection of phage resistant receptor mutants

Phage-resistant strains were generated using both liquid and solid culture methods:

#### Liquid culture method

Log-phase TK001 culture was diluted 100-fold, and 100 μL of high-titer phage suspension (>10^8^ PFU/mL, MOI > 10) was added. The mixture was incubated at 37°C with shaking at 40 rpm in a compact rocking incubator (TVS062CA, Advantec Toyo Co. Ltd., Tokyo, Japan). The emergence of resistant bacteria was confirmed by turbidity curves the following day. The culture was centrifuged (9000 rpm, 5 min, 4 °C), and the pellet was washed with 1× phosphate-buffered saline (PBS). The pellet was serially diluted in 1× PBS (1 mL), and 20 μL of each dilution was spotted onto soft agar containing 100 μL of high-titer phage. Eight phage-resistant *E. coli* colonies were isolated for each phage and purified by streak culture at least twice. Purified colonies were inoculated into 1 mL of LB, cultured for 11 hours, and stored at −80 °C.

#### Solid culture method

100 μL of overnight-cultured TK001 and 100 μL of high-titer phage suspension (>10^8^ PFU/mL) were added to soft agar and overlaid on agar plates. Four colonies per phage (one for ΦWec190 and three for ΦWec193) were isolated the following day and purified by streak culture at least twice. Purified colonies were processed and stored as described in the liquid culture method.

Phage adsorption assay was used to select target receptor mutants from the acquired phage-resistant strains. A mixture of overnight-cultured resistant strain (100 μL) and LB (890 μL) was combined with phage suspension (>10^7^ PFU/mL, 100 μL). Samples (10 μL) were taken at 0- and 10-min post-phage addition, mixed with 990 μL SM buffer (pre-added with 100 μL chloroform), and centrifuged (15,000 rpm, >2 min, 4 °C). The supernatant (100 μL) and overnight-cultured TK001 (100 μL) were mixed in soft agar and overlaid on agar plates. After overnight incubation, the unadsorbed ratio was calculated by dividing the number of plaques at 10 min by the number at 0 min. Strains with an adsorption rate (100 - unadsorbed ratio) below 20% were designated as target receptor mutants.

### Whole genome analysis of wild type strain

DNA was extracted from overnight static cultures of *E. coli* TK001 in LB medium using ISOPLANT II (Nippon Gene, 310-04151, Tokyo, Japan) according to the manufacturer’s instructions.

Sequencing was performed by BGI Genomics. Continuous genome sequencing was assembled using two different sequencing technologies. Shotgun sequencing was performed using DNBSEQ G400 (MGI) with a read length of 150 bp. Clean data was generated using SOAPnuke (BGI) to remove adapters and low-quality reads (parameters: “-n 0.01 −l 20 -q 0.4 --adaMis 3 --outQualSys 1”)(19). Long reads obtained by Oxford Nanopore Technologies were processed using porechop (ver. 0.2.4, default parameters). The processed shotgun and long-read data were assembled using SPAdes (ver. 3.15.5, parameters: “-k auto --cov-cutoff auto --careful -t 12”) for hybrid assembly (20). Annotation of the resulting contigs was performed using prokka (21). Basic statistical analysis of the contigs was conducted using seqkit (22).

### Whole genome analysis and mutation detection in mutant strains

DNA was extracted from selected target receptor mutant strains as described for the wild-type strain. BGI performed shotgun sequencing and read processing similarly to the wild-type strain. The mutant strain shotgun reads were mapped to the indexed wild-type sequence using bwa. Samtools was used to convert the mapping results to BAM format and sort them. The sorted BAM files were indexed using samtools. Guided assembly was performed using pilon (ver. 1.24, parameters: “--vcf --tracks –changes --verbose”) with the index files (23). Non-synonymous mutations were detected by comparing each resistant strain’s sequence obtained through guided assembly with the wild-type strain using nucdiff (24).

### Complementation experiments

To confirm that the identified mutations in phage-resistant strains were directly responsible for the resistance phenotypes, we performed complementation experiments. Wild-type genes corresponding to the mutated genes in resistant strains were cloned into the low-copy plasmid pACYC184. This plasmid was selected for its stability in TK001, and dual antibiotic resistance markers (tetracycline and chloramphenicol).

Each gene of interest, along with its native promoter region, was PCR-amplified from TK001 genomic DNA. The amplified fragments were designed to replace the tetracycline resistance gene in pACYC184 (Nippon Gene Co., Ltd., Tokyo, Japan) using In-Fusion cloning (Takara Bio Inc., Shiga, Japan). Care was taken to avoid including unnecessary adjacent genes when incorporating promoter regions. The resulting constructs were initially transformed into *E. coli* JM109 (Takara Bio Inc., Shiga, Japan), and positive transformants were selected on LB agar with chloramphenicol (25 μg/mL). For each gene, approximately 10 colonies were isolated and verified by PCR amplification of the inserted gene and by confirming loss of tetracycline resistance.

Verified plasmid constructs were then isolated and transformed into the corresponding phage-resistant mutant strains using a standard chemical competent cell protocol (see also supplementary method). For each complementation, three independent transformants were isolated and verified by PCR to confirm the presence of the complementing gene. The complemented strains were then tested for restored phage sensitivity using spot tests as described in the “Spot test using target receptor mutants” section. EoP values were calculated relative to the wild-type strain TK001 carrying the empty pACYC184 vector. Biological replicates were performed 3 times.

The complemented strains were then tested for restored phage sensitivity using the same spot test method as described in the “Spot test using target receptor mutants” section. EoP values were calculated relative to the wild-type strain TK001 carrying the empty pACYC184 vector. Biological replicates were performed 3 times.

For *waaV* mutations, which showed variable complementation results, we performed additional experiments by expressing the mutant variants of *waaV* in wild-type TK001 to assess whether the mutant proteins might interfere with wild-type function.

### Protein structure prediction and analysis

To understand the molecular basis of how the identified mutations affect protein function, we predicted protein structure using AlphaFold2 via Google Colaboratory (25)(26) or AlphaFold3 (27). For each mutated protein (WaaG, WaaV, WaaW, WaaT, WaaY, WecA, and ManB-1), we generated structural models of wild-type. The predictions were run with default parameters, and models were evaluated using the predicted local distance difference test (pLDDT) and predicted aligned error (pAE).

For WaaG, where crystal structures of homologs are available, predicted WaaG structure was superimposed on a previously solved crystal structure (PDB: 2IW1) to identify catalytic and substrate-binding sites. For WaaY and ManB1, the complex with ATP or GTP were predicted using AlphaFold3. For other proteins, potential binding sites were predicted using AutoDock4.2.6 (28) with UDP-glucose or UDP-galactose as ligands, as appropriate for each enzyme’s function.

Electrostatic surface potentials were calculated using the Adaptive Poisson-Boltzmann Solver (APBS) program. All structural analyses and visualizations were performed using PyMOL (version 3.1.4.1).

### Preparation of dried defatted bacterial cells

Dried defatted bacterial cells were prepared for LPS extraction. Bacterial cells were cultured in 2 L of medium and harvested by centrifugation (4,800 rpm, 15 min, 4°C) in four 500 mL centrifuge tubes. From *E. coli* cultures, approximately 20 g of wet cells were typically obtained. Cell pellets were washed by resuspension in 30 mL of 1× PBS and collected by centrifugation (4,800 rpm, 15 min, 4°C) in 50 mL tubes.

The cells were then defatted through sequential washing steps. First, cells were resuspended in 30 mL of 100% ethanol using a spatula, followed by centrifugation (4,800 rpm, 15 min, 4°C). This ethanol washing step was repeated until the cell pellet lost its sticky consistency (typically three times), with the supernatant turning yellow. The pellet was then washed once with 30 mL acetone, using gentle deceleration during centrifugation to prevent pellet disruption. Finally, cells were washed with 30 mL of hexane or diethyl ether.

The resulting pellet was dried under vacuum in a desiccator for 1-3 days. This process typically yielded 4-5 g of off-white to white dried defatted bacterial cells from *E. coli* cultures.

### LPS structure analysis

LPS structure analysis of wild type and target receptor mutant strains was performed using two methods: pathway analysis to predict the LPS synthesis pathway from retained genes, and sodium deoxycholate-polyacrylamide gel electrophoresis (DOC-PAGE) for LPS structure analysis.

#### Pathway analysis

KEGG Mapper was used to map retained genes to known synthesis pathways based on Kegg Orthology (KO) assignment results obtained using BlastKOALA (29),(30),(31).

#### Structural analysis

To obtain LPS from bacterial cells, dried defatted bacterial cells were first prepared. LPS extraction from dried defatted bacterial cells was performed using the hot phenol-water extraction method (32). Extracted LPS was visualized by DOC-PAGE followed by silver staining (33). Silver staining was performed using the Silver Stain MS Kit (Fujifilm Wako, 299-58901, Osaka, Japan), with a modified sodium periodate oxidation step between first and second fixative solutions (34). The gel was shaken in 10 times its volume of 1% w/v sodium periodate solution for 20 min, followed by three 5-min washes with MilliQ water. As control samples for the LPS structure of TK001, LPS extracted from the following strains was used: *Salmonella enterica* serovar Typhimurium (Sigma; catalogue no. L6511), *Escherichia coli* O111:B4 (Sigma; catalogue no. L4130), and various *S. enterica* serovars with different chemotypes (LPSs were extracted for bacteria of kindly provided by Dr. Hajime Ikigai). For presentation in Fig. 3(c), lanes from the original gel were rearranged for logical comparison without altering the relative positions or intensities of the bands. The original, unedited gel images were included in the supplementary materials (Fig. S11) for reference, ensuring transparency in the data presentation.

### Spot test using target receptor mutants

Each phage was serially diluted 10-fold up to 10^-7^ and spotted (1 μL) onto target receptor mutant lawns (35). The efficiency of plating (EoP) of the mutants relative to the wild type was calculated. Biological replicates were performed 3-7 times.

### Spot test using the Keio collection

For the five *E. coli* phages (ΦWec187, 189, 191, 193, and 196) capable of infecting K-12 strains(10), spot tests were conducted using the Keio collection (NBRP, Japan), a comprehensive gene knockout library (36),(37). The EoP of Keio collection strains relative to the wild-type BW25113 was calculated. The Keio collection strains used were limited to those predicted to be related to the target receptors of ΦWec187, 189, 191, 193, and 196 based on phage-resistant TK001 (Fig. S2). Biological replicates were performed 3 times.

### Whole genome construction of phages

Sequencing data were assembled using Velvet software (ver. 1.2.10) (38).

### Phylogenetic tree based on tail fibers

Whole genome information of each phage was annotated using RAST, and tail fiber sequences were extracted (39). Additionally, tail fibers of the closest relatives were obtained by BLAST search of each phage’s whole genome, and similar sequences were extracted from the whole genomes of phages drawing these sequences. The obtained sequences were translated into amino acids, and for each phage, the sequences were artificially combined into a single sequence. A phylogenetic tree was constructed using MEGA X with the Maximum-likelihood method and 1,000 bootstrap replicates (40).

### Whole genome phylogenetic tree and similarity

The whole genome phylogenetic tree was created using Viptree (41). Furthermore, phylogenetic analysis of the phages was conducted using vConTACT2 (42) with the reference genome of 3503 phage (ProkaryoticViralRefSeq201) from vConTACT2, and visualized using Cytospace v3.10.1 (43). The Average Nucleotide Identity (ANI) was calculated using pyani (44). The parameters used were: pyani: anib and ANI calculator: minimum identity 0%, with other parameters set to default.

### Phage grouping based on physiological characteristics, tail fibers, and whole genomes

Phages were grouped based on their physiological characteristics as previously reported. To improve the robustness of our previously proposed clustering method, we optimized the bacterial panel for host range evaluation by analyzing infection patterns of 58 phages (including phages previously reported in our previous study (15)) against 41 *Enterobacteriaceae* strains (Table S4). Using a greedy algorithm, we selected 16 strains that collectively maximized the discriminatory power while minimizing the number of strains required.

For data analysis, Gower distance was implemented instead of Euclidean distance to appropriately handle the mixed nature of our dataset, which contained both qualitative infection data (binary host range patterns) and quantitative measurements (continuous physiological parameters).

To objectively determine the optimal number of clusters, silhouette coefficient analysis was performed for all three grouping methods (physiological characteristics, tail fiber phylogenetics, and whole genome phylogenetics). For each potential number of clusters k (ranging from 2 to 9), we calculated the average silhouette coefficient across all phages in the dataset. The silhouette coefficient for each phage measures how similar it is to its own cluster compared to other clusters, with values ranging from −1 to 1. Higher values indicate better-defined clusters. The optimal number of clusters for each grouping method was determined as the value of k that maximized the average silhouette coefficient.

These analyses were performed using R (version 4.0.3) with the “cluster” and “factoextra” packages for silhouette calculations. Hierarchical clustering was performed using the ‘complete’ linkage method as previously reported (10).

### Comparison of similarity between methods

Data analysis was performed using Python 3.9.12, with major packages including pandas (1.4.2), numpy (1.22.3), and scikit-learn (1.0.2). The similarity between different grouping methods and with target receptor-based grouping was evaluated using Adjusted Rand Index (ARI). Label normalization was performed in the similarity evaluation to improve the accuracy of comparison between methods. These analyses were executed on Linux.

## Data Availability

Phage genome sequences used in this study were deposited in the DNA Data Bank of Japan (DDBJ) with the following accession numbers: ΦWec172-ΦWec196 (LC739530-LC739542), ΦWec270 (DRX534190), and ΦWec272 (DRX534192). The T5 bacteriophage genome sequence was obtained from GenBank (accession number AY543070.1). All other data supporting the findings of this study are included within the manuscript and its supplementary information files.

## Acknowledgements

We extend our sincere gratitude to Midori Ishikawa and Hirohito Wakashima for providing the LPS samples used as controls in our DOC-PAGE analysis. Their contributions were crucial for our comparative study of *E. coli* LPS structures. Specifically, we acknowledge: Midori Ishikawa for isolating LPS from *S. enterica* serovar Typhimurium TV119 (Ra chemotype) and *S. enterica* serovar Typhimurium SL733 (Rb_1_ chemotype). Hirohito Wakashima for isolating LPS from *S. enterica* serovar Typhimurium SL1034 (semi-rough chemotype), *S. enterica* serovar Minnesota SF1119 (RcP^-^ chemotype), *S. enterica* serovar Minnesota SF1121 (Rd_1_P^-^ chemotype), *S. enterica* serovar Minnesota SF1118 (Rd_2_ chemotype), and *S. enterica* serovar Typhimurium TA2168 (Re chemotype). In addition to the previous acknowledgement of Matthew Imanaka for English language editing on the initial version, we would like to thank Claude 3.7 Sonnet (Anthropic) for its assistance in improving the English grammar and style of this revised manuscript.

