## Supplemental Material for "From phenotype to receptor: validating physiological clustering of *Escherichia coli* phages through comprehensive receptor analysis"

**Contents**

**Supplementary Methods … p2 - 6**

**Supplementary Results and Discussion (also Fig. S3 ~ S9) … p7 - 20**

**Supplementary Figures S1&2 … p21 - 22**

**Supplementary Tables … p23 - 26**

**References … p27**

**Supplementary Methods**

**Preparation of chemically competent *E. coli* cells**

This protocol describes the preparation of chemically competent *E. coli* cells using the calcium chloride method. The procedure renders bacterial cells permeable to plasmid DNA, enabling efficient transformation for complementation experiments.

***Materials and Reagents***

- *E. coli* glycerol stock
- LB agar plates
- LB liquid medium
- 100 mM CaCl₂ (pre-chilled to 4°C)
- 100 mM MgCl₂ (pre-chilled to 4°C)
- 100% glycerol
- 85 mM CaCl₂/15% glycerol solution (prepared fresh)
- Liquid nitrogen
- Sterile 5 mL and 200 μL pipette tips
- 50 mL conical centrifuge tubes
- 1.5 mL microcentrifuge tubes
- Ice bath

**Protocol**

1. Streak the *E. coli* glycerol stock onto an LB agar plate and incubate overnight at 37°C.
2. Inoculate a single colony into 2 mL of LB medium and incubate overnight at 37°C with shaking (120 rpm).
3. Day of preparation:

- Pre-chill LB medium (50 mL in a flask), 100 mM CaCl₂ (15 mL), 100 mM MgCl₂ (25 mL), and 100% glycerol at 4°C
- Prepare 5 mL of 85 mM CaCl₂/15% glycerol solution in a centrifuge tube
- Pre-chill pipette tips and tubes at -20°C

4. Inoculate 500 μL of the overnight culture into 50 mL of pre-chilled LB medium in a flask. Incubate at 37°C with shaking until the OD₆₀₀ reaches approximately 0.5 (approximately 2 hours).
5. Monitor the OD₆₀₀ every 20 minutes once the culture begins to become turbid. Simultaneously, prepare an ice bath and set the centrifuge to 4°C.
6. When the OD₆₀₀ reaches approximately 0.5, transfer the culture to a 50 mL centrifuge tube and place on ice for 10 minutes to rapidly cool the cells.
7. Centrifuge at 845 G for 10 minutes at 4°C.
8. Discard the supernatant by gentle decantation and add cold 100 mM MgCl₂ to approximately 15 mL mark on the tube.
9. Gently resuspend the pellet while keeping the tube on ice.
10. Centrifuge at 587 G for 5 minutes at 4°C with maximum deceleration time.
11. Discard the supernatant carefully as the pellet will be fragile. Add cold 100 mM CaCl₂ to approximately the 20 mL mark on the tube.
12. Gently resuspend the pellet while keeping the tube on ice.
13. Incubate the cell suspension on ice for 30-60 minutes.
14. Centrifuge at 587 G for 5 minutes at 4°C with maximum deceleration time.
15. Carefully discard the supernatant and gently add 5 mL of pre-chilled 85 mM CaCl₂/15% glycerol solution using a 5 mL pipette.
16. Gently resuspend the pellet while keeping the tube on ice until no visible clumps remain.
17. Aliquot 100 μL of the cell suspension into pre-chilled 1.5 mL microcentrifuge tubes.
18. Flash-freeze the tubes in liquid nitrogen for at least 1 minute.
19. Transfer the frozen tubes to a -80°C freezer for long-term storage.

Note: All operations should be performed under aseptic conditions. The competent cells prepared by this method were used for transformation of plasmid constructs in the complementation experiments described in the main text.

**R Scripts for Phage Clustering Analysis**

The following R code was used to perform clustering analyses based on physiological characteristics, whole genome phylogeny, and tail fiber phylogeny. For each analysis method, silhouette coefficient analysis was implemented to objectively determine the optimal number of clusters.

For whole genome phylogeny, a Newick-format phylogenetic tree file (.newick or .nwk extension) generated by the VipTree software suite was used. For tail fiber phylogeny, a Newick-format tree file (.ntw or .newick extension) derived from MEGA X analysis of aligned tail fiber protein sequences was utilized.

| Clustering Based on Physiological Characteristics | Clustering Based on Whole Genome Phylogeny | Clustering Based on Tail Fiber Phylogeny |
| --- | --- | --- |
| # Required packages  library(cluster)  library(factoextra)  library(openxlsx)  library(ggplot2)  library(dendextend)  # Load data  data <- read.xlsx("phage_physiological_data.xlsx"")  # Convert time measurements to minutes if applicable  convert_decimal_to_minutes <- function(decimal_time) {  if (is.na(decimal_time)) return(NA)  return(decimal_time * 24 * 60)  }  # Define variable types  binary_vars <- c("MG1655", "BL21", "ESBL953", "NBRC102203", "ESBL983",  "TOP10F", "ESBL946", "ESBL1054", "ESBL1013", "SP")  quant_vars <- c("onset_minutes", "duration_minutes", "adsorption_constant", "burst_size", "phage_yield")  # Create analysis dataframe  analysis_data <- data[, c(binary_vars, quant_vars)]  rownames(analysis_data) <- data$phage  # Calculate Gower distance  gower_dist <- daisy(analysis_data,  metric = "gower",  type = list(asymm = binary_vars,  symm = character(0),  numeric = quant_vars))  # Hierarchical clustering  hc <- hclust(gower_dist, method = "ward.D2")  # Calculate silhouette scores for different numbers of clusters  max_k <- min(9, nrow(analysis_data) - 1)  sil_width <- numeric(max_k)  sil_width[1] <- NA # Silhouette not defined for k=1  for (i in 2:max_k) {  clusters_i <- cutree(hc, k = i)  sil_obj <- silhouette(clusters_i, gower_dist)  sil_width[i] <- mean(sil_obj[, "sil_width"])  }  # Find optimal number of clusters  best_k <- which.max(sil_width[-1]) + 1  cat("Optimal number of clusters based on silhouette method: ", best_k)  # Get the final clusters  clusters <- cutree(hc, k = best_k) | # Required packages  library(ape)  library(cluster)  library(factoextra)  # Load the phylogenetic tree from Newick file  tree_file <- "whole_genome_tree.newick"  tree <- read.tree(tree_file)  # Extract distance matrix from the tree  dist_matrix <- cophenetic.phylo(tree)  # Calculate silhouette scores for different numbers of clusters  max_k <- min(9, nrow(dist_matrix) - 1)  sil_width <- numeric(max_k)  sil_width[1] <- NA # Silhouette not defined for k=1  for (i in 2:max_k) {  # Perform hierarchical clustering on the distance matrix  hc <- hclust(as.dist(dist_matrix), method = "ward.D2")  clusters_i <- cutree(hc, k = i)    # Calculate silhouette coefficient  sil_obj <- silhouette(clusters_i, as.dist(dist_matrix))  sil_width[i] <- mean(sil_obj[, "sil_width"])  }  # Find optimal number of clusters  best_k <- which.max(sil_width[-1]) + 1  cat("Optimal number of clusters based on silhouette method: ", best_k)  # Get the final clusters  hc <- hclust(as.dist(dist_matrix), method = "ward.D2")  clusters <- cutree(hc, k = best_k) | # Required packages  library(ape)  library(cluster)  library(factoextra)  # Load the phylogenetic tree from Newick file  newick_file <- "tail_fiber_tree.newick"  tree <- read.tree(newick_file)  # Calculate cophenetic distance matrix from phylogenetic tree  cophenetic_matrix <- cophenetic(tree)  # Perform hierarchical clustering  hc <- hclust(as.dist(cophenetic_matrix), method = "ward.D2")  # Silhouette analysis to determine optimal cluster number  silhouette_scores <- c()  for (k in 2:9) {  cluster_assignment <- cutree(hc, k = k)  sil <- silhouette(cluster_assignment, as.dist(cophenetic_matrix))  silhouette_scores[k-1] <- mean(sil[, "sil_width"])  }  # Identify optimal cluster number  optimal_k <- which.max(silhouette_scores) + 1  cat("Optimal number of clusters based on silhouette method: ", optimal_k)  # Get the final clusters  clusters <- cutree(hc, k = optimal_k) |

**Supplementary Results, Discussion, and Fig. S3 ~ S9**

**Structural prediction analysis of mutated proteins**

**WaaG**

The binding site of UDP-Glucose was analyzed based on a previously crystallized structure available in the literature: "Insights into the Synthesis of Lipopolysaccharide and Antibiotics through the Structures of Two Retaining Glycosyltransferases from Family GT4" (1) and PDB: 2IW1 (<https://www.rcsb.org/structure/2IW1>).

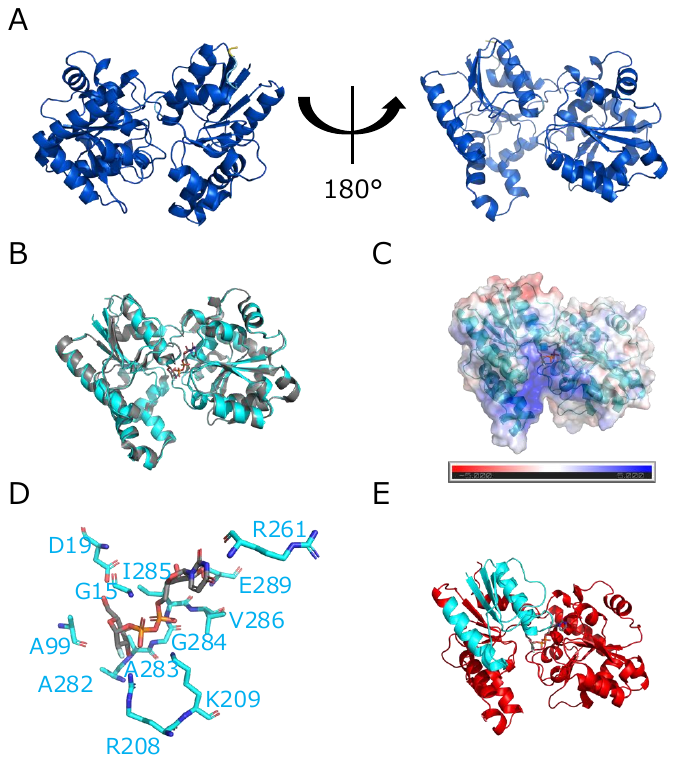

Fig. S3 Structure analysis of WaaG model (A) The structure of WaaG was predicted by AlphaFold2. The pLDDT scores were displayed in blue for >90, cyan for >70, yellow for >50, and red for <50. (B) AlphaFold2 model of WaaG (cyan) was superimposed on the crystal structure 2IW1, which is bounded with UDP-2-deoxy-2-fluoro glucose (gray). (C) The electrostatic surface of the WaaG model was visualized. Charges on the WaaG surface are colored according to their electrostatic properties. The scale bar indicates electrostatic property values ranging from -5.0 kT/e (red) to 5.0 kT/e (blue). (D) The residues interacting with the substrate were shown. (E) The truncated region was highlighted in red.

The pLDDT score was above 90 for nearly all residues. The only residues with slightly lower pLDDT scores were Phe-13, Gly-14, Gly-370, Gly-371, Leu-372 (cyan), and Asp-373, Gly-374 (yellow) (Figure S3A). The structural alignment between the WaaG model and the crystcrystal structure (2IW1) showed excellent agreement (Figure S3B). The sequence alignment between WaaG and the crystallized homolog showed 90.1% identity (338/375 residues) and 93.9% similarity (352/375 residues), further supporting the validity of the structural model. The electrostatic surface potential analysis revealed a region of positive charge (blue) that likely interacts with the negatively charged phosphate groups of UDP-Glucose (Figure S3C). It was suggested that negatively charged LPS may bind to a positively charged pocket near the substrate-binding site. Based on the structural analysis, residues Gly-15, Asp-19, Ala-99, Arg-208, Lys-209, Arg-261, Ala-282, Ala-283, Gly-284, Ile-285, Val-286, and Glu-289 are likely involved in UDP-Glucose binding (Figure S3D). These residues correspond to those identified in the reference crystal structure. The nonsense mutation in the phage-resistant strain would result in a truncated protein missing almost all of these binding residues, which is consistent with the observed complete loss of enzymatic function (Figure S3E).

**WaaV**

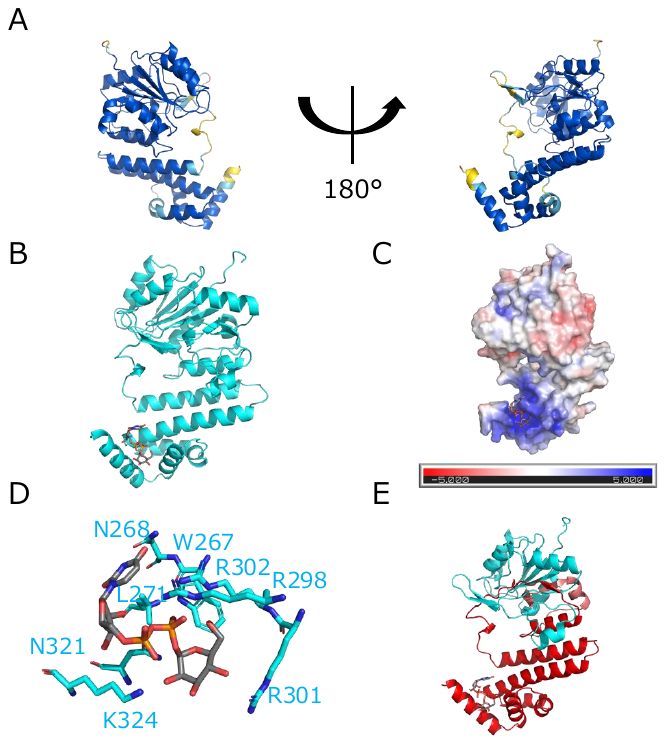

Fig. S4 Structure analysis of WaaV model (A) The structure of WaaV was predicted by AlphaFold2. The pLDDT scores were displayed in blue for >90, cyan for >70, yellow for >50, and red for <50. (B) UDP-Glucose (gray) binding WaaV model (cyan) was predicted using Autodock4. (C) The electrostatic surface of the WaaV model was visualized. Charges on the WaaV surface are colored according to their electrostatic properties. The scale bar indicates electrostatic property values ranging from -5.0 kT/e (red) to 5.0 kT/e (blue). (D) The residues interacting with the substrate were shown. (E) The truncated region was highlighted in salmon (158) and red (181).

Approximately 90% of the residues had pLDDT scores above 90, indicating high confidence in the predicted structure. Regions with slightly lower confidence were primarily located in terminal regions and flexible loops (Figure S4A). UDP-Glucose binding to the WaaV model was predicted using Autodock4 (Figure S4B), and electrostatic analysis revealed a positively charged pocket (blue) that likely interacts with the negatively charged phosphate groups of UDP-Glucose (Figure S4C). The structural analysis identified Trp-267, Asn-268, Leu-271, Arg-298, Arg-301, Arg-302, Asn-321, and Lys-324 as potential residues involved in substrate binding (Figure S4D). The frameshift mutation at position 158 and 181 would result in a truncated protein missing all of these binding residues (Figure S4E), which is consistent with the absence of O-antigen in the DOC-PAGE analysis of the mutant strains.

**WaaW**

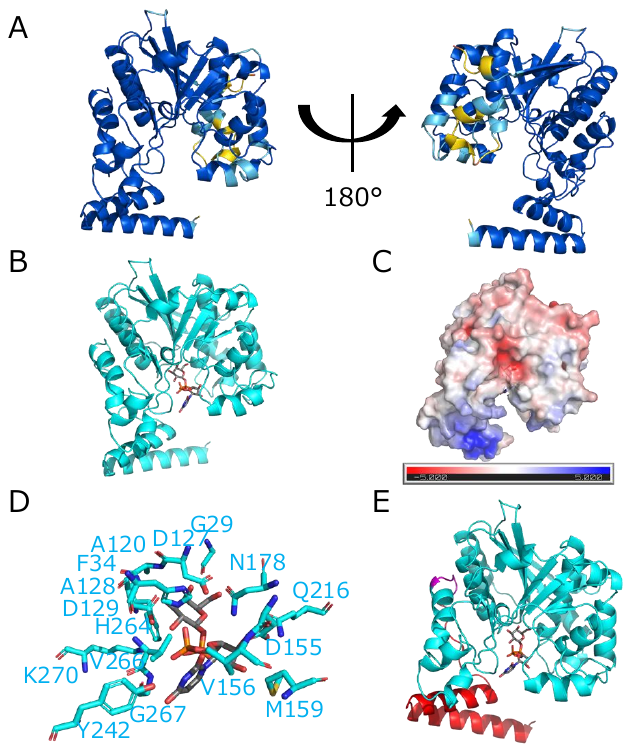

Fig. S5 Structure analysis of WaaW model (A) The structure of WaaW was predicted by AlphaFold2. The pLDDT scores were displayed in blue for >90, cyan for >70, yellow for >50, and red for <50. (B) UDP-Galactose (gray) binding WaaW model (cyan) was predicted using Autodock4. (C) The electrostatic surface of the WaaW model was visualized. Charges on the WaaW surface are colored according to their electrostatic properties. The scale bar indicates electrostatic property values ranging from -5.0 kT/e (red) to 5.0 kT/e (blue). (D) The residues interacting with the substrate were shown. (E) The mutated region was highlighted in magenta. The truncated region was highlighted in red.

Approximately 83% of residues had pLDDT scores above 90, with the majority of lower-confidence regions located in terminal regions and flexible loops (Figure S5A). UDP-Glucose binding of WaaW model was predicted using Autodock4 (Figure S5B), and the binding pocket region was predicted to interact with UDP-Galactose (Figure S5C). Gly-29, Phe-34, Asp-127, Ala-128, Asp-129, Asp-155, Val-156, Met-159, Asn-178, Gln-216, Tyr-242, His-264, Val-266, Gly-267, and Lys-270 were identified as potential residues involved in substrate binding (Figure S5D). The frameshift mutation at position 290 would not directly affect these active site residues, as they remain intact in the truncated protein (Figure S5E). However, the mutation disrupts a C-terminal region rich in basic amino acids that likely participates in interactions with the negatively charged phosphate groups of LPS. This explains why the mutant shows altered LPS structure despite retaining most of the catalytic residues.

**WaaT**

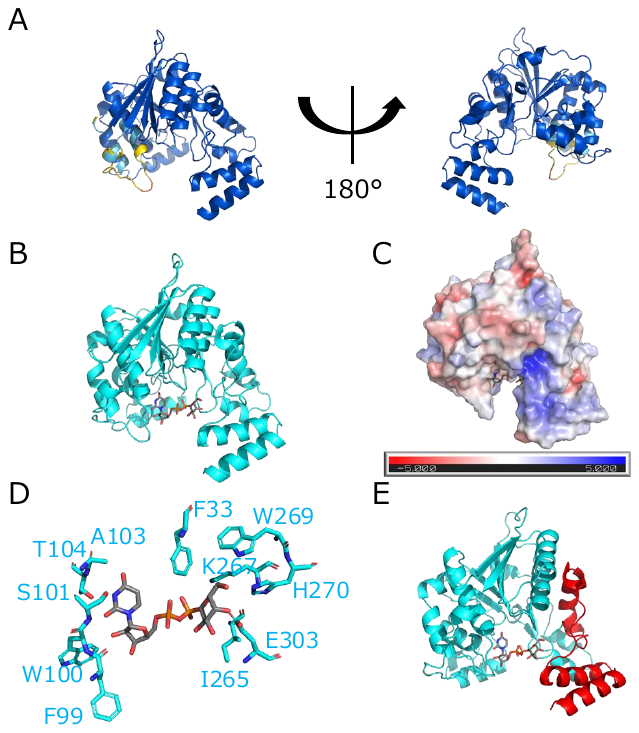

Fig. S6 Structure analysis of WaaT model (A) The structure of WaaT was predicted by AlphaFold2. The pLDDT scores were displayed in blue for >90, cyan for >70, yellow for >50, and red for <50. (B) UDP-Galactose (gray) binding WaaT model (cyan) was predicted using Autodock4. (C) The electrostatic surface of the WaaT model was visualized. Charges on the WaaT surface are colored according to their electrostatic properties. The scale bar indicates electrostatic property values ranging from -5.0 kT/e (red) to 5.0 kT/e (blue). (D) The residues interacting with the substrate were shown. (E) The truncated region was highlighted in red.

Approximately 90% of residues had pLDDT scores above 90, with only a few regions showing lower confidence scores (Figure S6A). UDP-Glucose binding of WaaT model was predicted using Autodock4 (Figure S6B), and the binding pocket region was predicted to interact with UDP-Galactose (Figure S6C). Phe-33, Phe-99, Trp-100, Ser-101, Ala-103, Thr-104, Ile-265, Lys-267, Trp-269, His-270, and Glu-303 were identified as potential residues involved in substrate binding (Figure S6D). The frameshift mutation after position 269 would result in the loss of some of these binding residues, including His-270 and Glu-303. Notably, WaaT and WaaW share approximately 58% sequence similarity, and both exhibit regions rich in basic amino acids that are disrupted by the mutations (Figure S6E).

**WaaY**

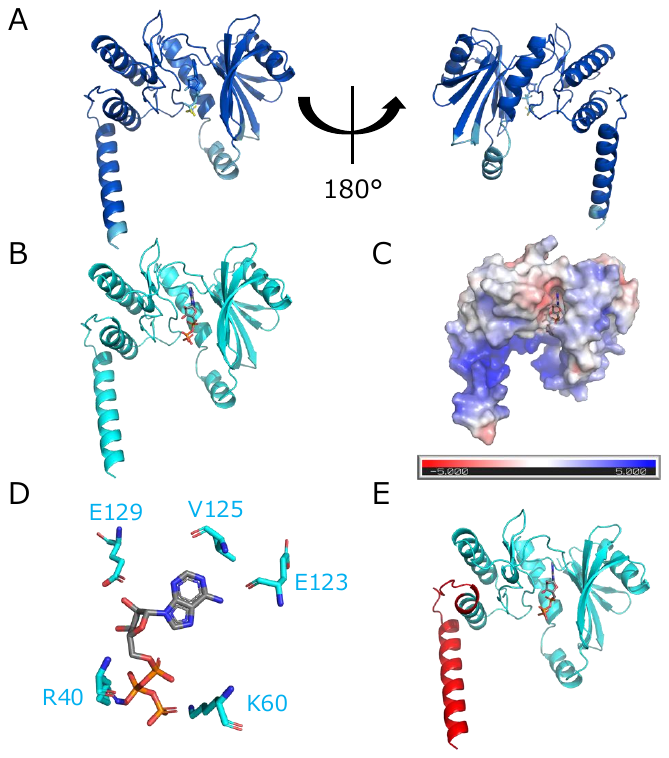

Fig. S7 Structure analysis of WaaY model (A) The structure of WaaY-ATP complex was predicted by AlphaFold3. The pLDDT scores were displayed in blue for >90, cyan for >70, yellow for >50, and red for <50. (B) ATP (gray) binding WaaY model (cyan) was shown. (C) The electrostatic surface of the WaaY model was visualized. Charges on the WaaY surface are colored according to their electrostatic properties. The scale bar indicates electrostatic property values ranging from -5.0 kT/e (red) to 5.0 kT/e (blue). (D) The residues interacting with the substrate were shown. (E) The truncated region was highlighted in red.

Approximately 85% of residues had pLDDT scores above 90, indicating high confidence in the majority of the structure (Figure S7A). ATP binding of WaaY model was predicted using AlphaFold3 (Figure S7B), and the binding pocket region was predicted to interact with ATP (Figure S7C). Arg-40, Lys-60, Glu-123, Val-125 and Glu-129 were identified as potential residues involved in substrate binding (Figure S7D). The frameshift mutation at position 169 would not directly affect these active site residues, as they remain intact in the truncated protein (Figure S7E). However, the mutation disrupts a C-terminal region rich in basic amino acids that likely participates in interactions with the negatively charged phosphate groups of LPS. This explains why the mutant shows altered LPS structure despite retaining most of the catalytic residues. The nonsense mutation at position 196 would result in a truncated protein missing a C-terminal region rich in basic amino acids, explaining why phages requiring phosphorylated heptose could not infect these mutants despite no visible change in overall LPS length in DOC-PAGE.

**WecA**

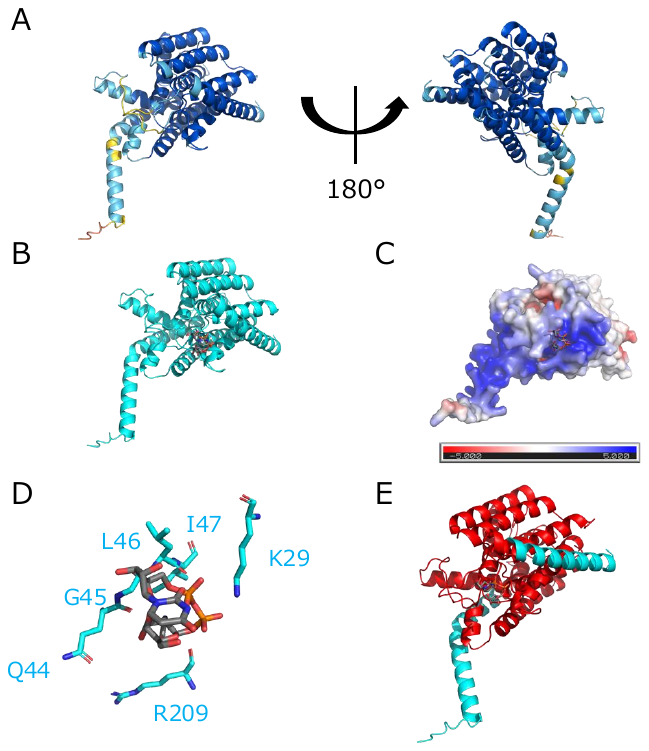

Fig. S8 Structure analysis of WecA model (A) The structure of WecA was predicted by AlphaFold2. The pLDDT scores were displayed in blue for >90, cyan for >70, yellow for >50, and red for <50. (B) UDP-GlcNAc (gray) binding WecA model (cyan) was predicted using Autodock4. (C) The electrostatic surface of the WecA model was visualized. Charges on the WecA surface are colored according to their electrostatic properties. The scale bar indicates electrostatic property values ranging from -5.0 kT/e (red) to 5.0 kT/e (blue). (D) The residues interacting with the substrate were shown. (E) The truncated region was highlighted in red.

Approximately 72% of residues had pLDDT scores above 90, with some regions showing moderate to low confidence (Figure S8A). UDP**-**GlcNAc binding of WecA model was predicted using Autodock4 (Figure S8B), and the electrostatic analysis showed a positively charged pocket (blue) that likely interacts with the negatively charged phosphate groups of UDP**-**GlcNAc (Figure S8C). Lys-29, Gln-44, Gly-45, Leu-46, Ile-47 and Arg-209 were identified as potential residues involved in substrate binding (Figure S8D). The nonsense mutation in the phage-resistant strain would result in a truncated protein missing almost all of these binding residues, which is consistent with the observed complete loss of enzymatic function (Figure S8E).

**ManB-1**

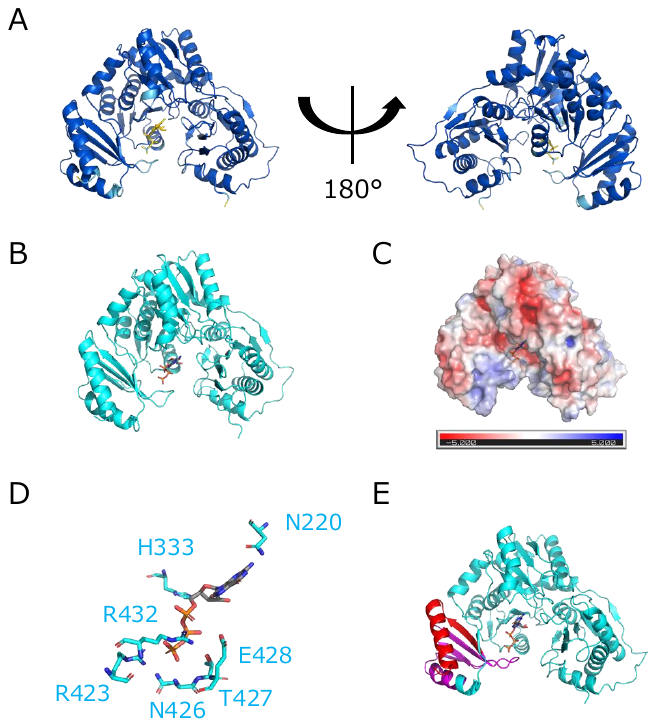

Fig. S9 Structure analysis of ManB-1 model (A) The structure of ManB-1-GTP complex was predicted by AlphaFold3. The pLDDT scores were displayed in blue for >90, cyan for >70, yellow for >50, and red for <50. (B) GTP (gray) binding ManB-1 model (cyan) was shown. (C) The electrostatic surface of the ManB-1 model was visualized. Charges on the ManB-1 surface are colored according to their electrostatic properties. The scale bar indicates electrostatic property values ranging from -5.0 kT/e (red) to 5.0 kT/e (blue). (D) The mutated region was highlighted in magenta. The truncated region was highlighted in red.

Approximately 94% of residues had pLDDT scores above 90, indicating very high confidence in the predicted structure. (Figure S9A). GTP binding of ManB-1 model was predicted using AlphaFold3 (Figure S9B), and the binding pocket region was predicted to interact with GTP (Figure S9C). Asn-220, His-330, Arg-423, Asn-426, Thr-427, Glu-428 and Arg-432 were identified as potential residues involved in substrate binding (Figure S9D). The frameshift mutation at position 390 would result in a mutated and truncated protein missing almost all of these binding residues, which is consistent with the observed complete loss of enzymatic function (Figure S9E).

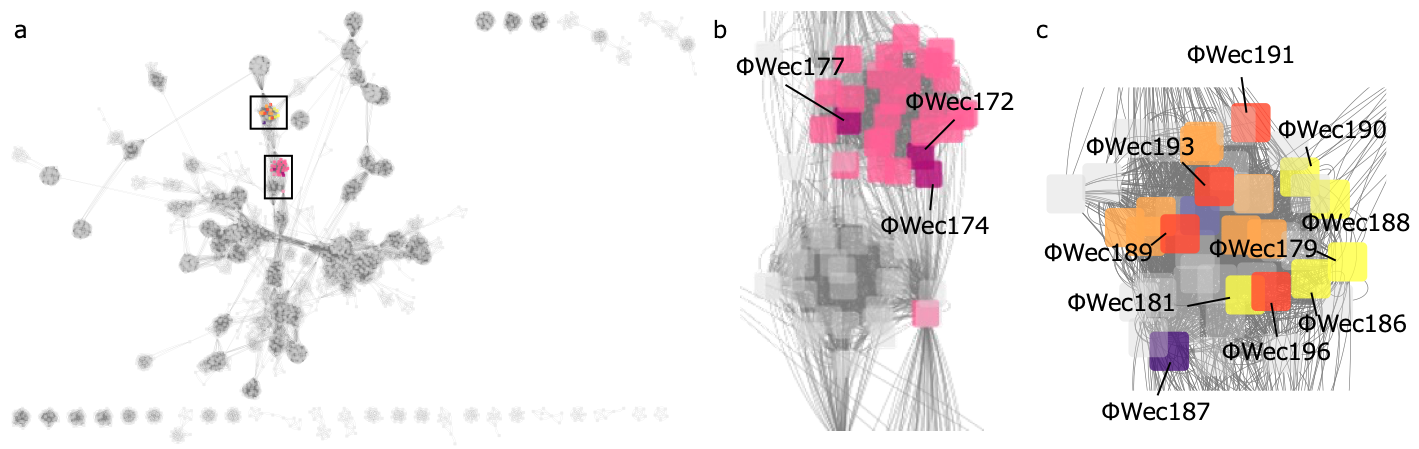

**Fig. S1 Visualized physiological relationship.** The genomes of the 13 phages used in this study and the genomes of 3503 reference phages (Prokaryotic Viral RefSeq201) from vConTACT2 were clustered based on shared proteins and analyzed as a network. The distance between phages was measured using vConTACT2, and the network was visualized using Cytoscape. (a) Overview of the entire network showing viral clusters. (b) Enlarged view of the cluster containing ΦWec172, 174, and 177. All phages in this viral cluster belong to various genera within *Ounavirinae*. (c) Enlarged view showing ΦWec179, 181, 186, 187, 188, and 190, which belong to or are closely related to viral clusters containing phages exclusively from various genera within *Stephanstirmvirinae* (2). Additionally, ΦWec189, 191, 193, and 196 belong to a viral cluster containing only *Vequintavirus* phages from *Vequintavirinae*.

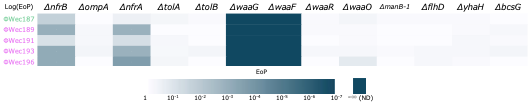

**Fig. S2 Heatmap of infectivity of K-12-infecting phages against Keio collection strains**

Five phages capable of infecting *E. coli* K-12 were tested against single-gene knockout strains from the Keio collection. The heatmap displays EoP values calculated relative to the wild-type BW25113 strain. White indicates no reduction in infectivity (EoP = 1), and progressively darker cyan shades represent decreasing infectivity levels, with dark cyan signifying complete loss of infectivity (not detected, ND). Results confirm the essential roles of specific genes in phage reception across different *E. coli* genetic backgrounds.

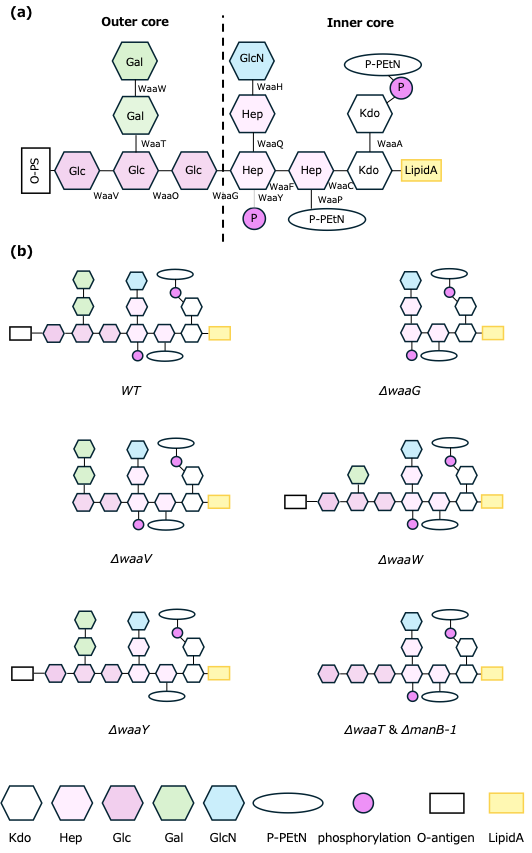

**Fig. S10 Prediction of LPS synthesis pathway in *Escherichia coli* TK001 strain by KEGG mapping**

(a) Structure of R1-type R-core and related synthesis enzymes. (b) Predicted LPS structures of LPS-related gene mutants.

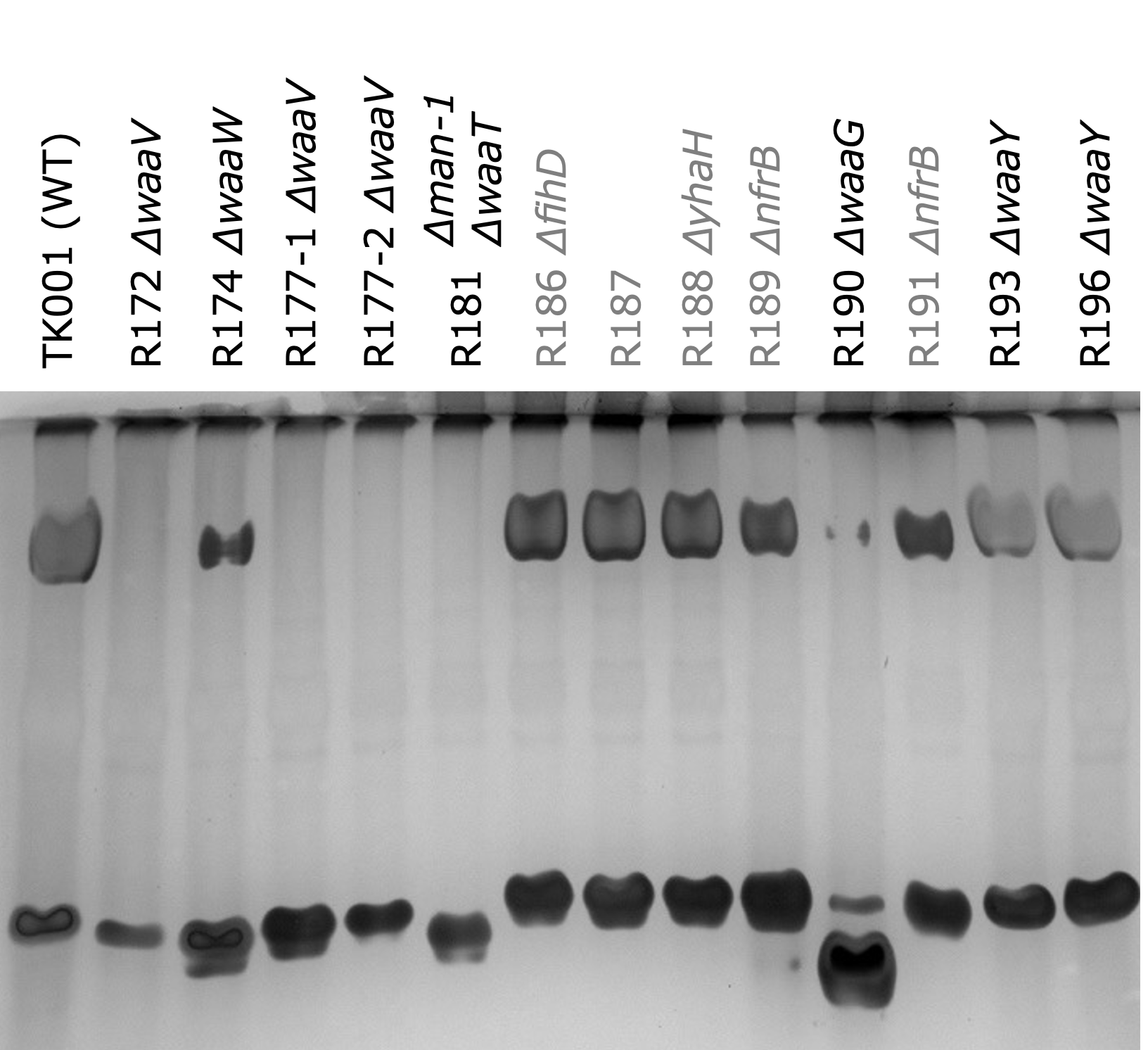

**Fig. S11 Original unedited DOC-PAGE gel images of LPS from wild-type TK001 and phage-resistant mutant strains**

Original, unedited DOC-PAGE gel image showing LPS from wild-type TK001 and all phage-resistant mutant strains. All samples were run simultaneously on the same gel under identical conditions, ensuring accurate size comparisons between samples. In Fig. 1(c) of the main text, lanes from this gel have been rearranged for ease of viewing and logical comparison, but no other adjustments were made to the image that would affect the relative positions or intensities of the bands.

**Table S1 ANI of phages in this study**
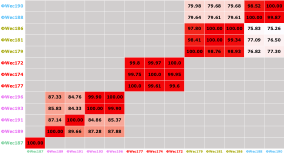

**Table S2 ANI between ΦWec172, 174, 177 and registered *Ounavirinae* phages**

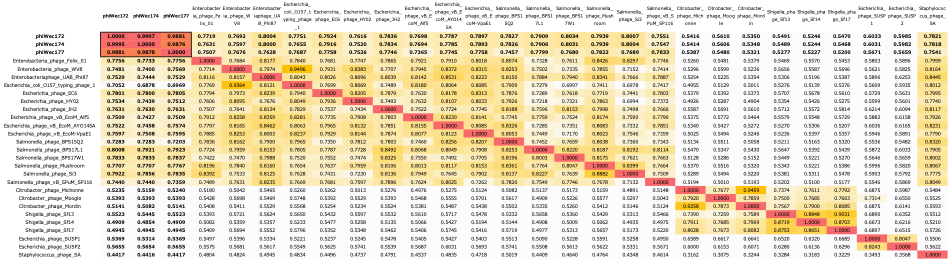

The table shows ΦWec172, 174, and 177 in the upper left. Maximum ANI values with registered phages range between 70% and 80%, suggesting these phages represent new species within the *Felixounavirus* genus of *Ounavirinae*, considering the established thresholds of 70% for genus demarcation and 95% for species demarcation. For detailed data, refer to the supplementary information Excel file.

**Table S3 ANI between ΦWec189, 191, 193, 196 and registered *Vequintavirus* phages**

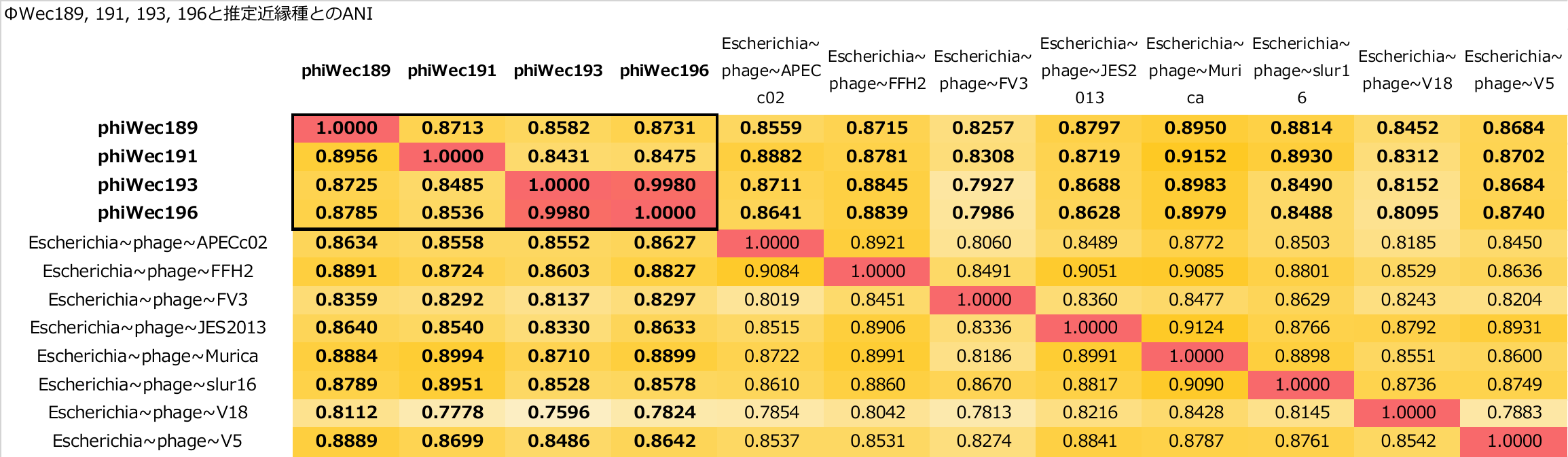

The table shows ΦWec189, 191, 193, and 196 in the upper left. Maximum ANI values with registered phages range between 70% and 90%, suggesting these phages represent new species within the *Vequintavirus* genus, considering the established thresholds of 70% for genus demarcation and 95% for species demarcation. For detailed data, refer to the supplementary information Excel file.

**Table S4 Bacterial strains tested for optimization of host range assessment panel**

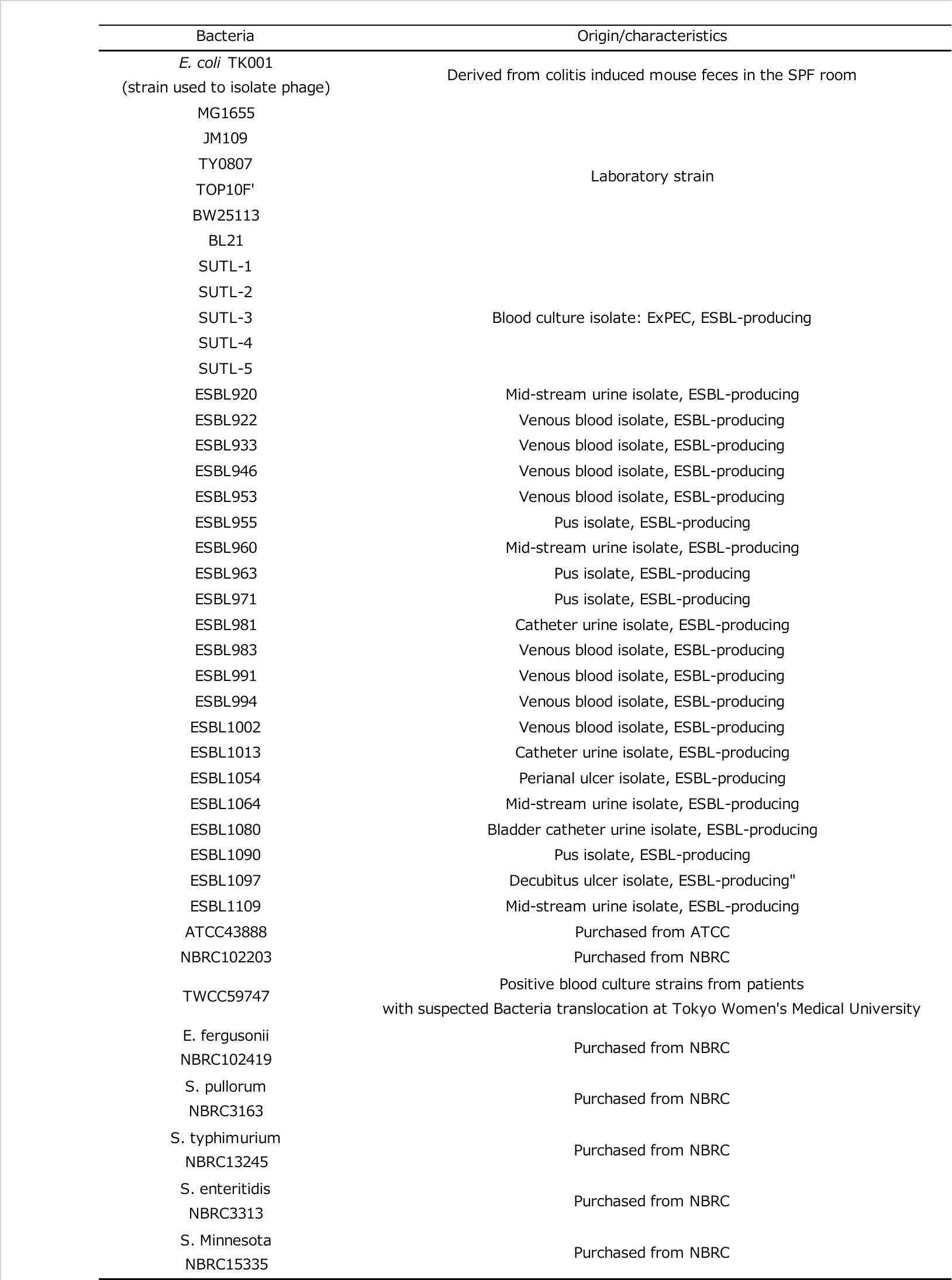

**Table S5 Infection matrix showing host range patterns of phages against bacterial strains**

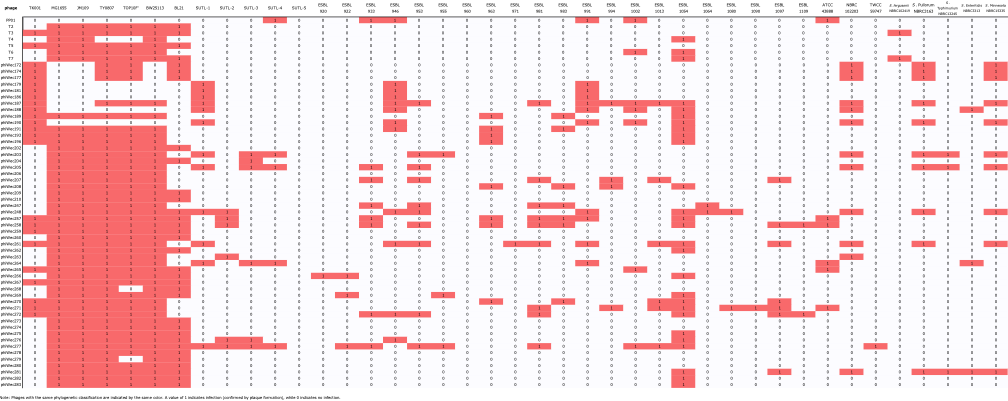

A value of 1 indicates infection (confirmed by plaque formation), while 0 indicates no infection.
